# Investigating the Impact of Omental Adipocytes on Ovarian Cancer: Insights from a 3D in-silico model

**DOI:** 10.1101/2025.07.03.662794

**Authors:** S. Oliver, M. Williams, D. Howard, D. Gonzalez, G. Powathil

## Abstract

Ovarian cancer is responsible for the most deaths of all gynaecological cancers in the Western world [1]. The symptoms of ovarian cancer are typically subtle and alike to less lethal diseases found more prevalently in the population, frequently resulting in late diagnoses and advanced tumour stages upon treatment initiation [2]. While surgery and platinum-based treatments can be curative, ovarian cancers found at the latter, metastasised stages are likely to be recurrent and more tailored towards palliative care [3]. Metastasised ovarian cancer spreads to surrounding organs and tissues such as the greater omentum [4], a large fat pad composed of adipose tissue stretching from the stomach and hanging over the intestines. The location of this is key in its role towards ovarian cancer and its progression [5]. In this study, we develop a mathematical model to investigate the role that adipocytes found in adipose tissue can have in ovarian cancer progression. Observations of biological experiments from two cell lines create foundations to build a multiscale agent-based model in a Physicell framework. The impact of the adipose derived media concentration, treatment dosage, and initial tumour size are explored to find how these conditions affect the spatio-temporal dynamics of cancer tumours.

## 2 Introduction

Ovarian cancer is the most lethal gynaecological malignancy worldwide [6]. Once regarded as a single entity, it is now recognised as a heterogeneous group of tumours with distinct origins, molecular features, clinical presentations, treatment responses and prognoses [7]. Risk factors include age, BRCA1/2 mutations, family history, endometriosis, hormone-replacement therapy, nulliparity, and, crucially, obesity [8]. Epidemiological evidence highlights excess adiposity, often measured by body-mass index (BMI) or visceral fat, as a modifiable driver of both carcinogenesis and poorer survival. A 2007 meta-analysis of 28 studies linked obesity to increased ovarian-cancer risk in 24, across all histological subtypes [9], while subsequent analyses report reduced overall survival, including in high-grade serous disease [10]. Collectively, these findings suggest that obesity and increased adiposity may not only drive carcinogenesis but also accelerate disease progression and metastatic potential. Clarifying how excess adiposity worsens ovarian-cancer outcomes therefore necessitates a closer examination of adipose tissue and its contribution to the ovarian tumour cancer microenvironment.

Adipose tissue comprises mature adipocytes plus a stromal vascular fraction (SVF) rich in stem cells, pre-adipocytes, macrophages and fibroblasts [11, 12]. Adipocytes originate from precursor cells and dominate tissue volume; each containing a large triglyceride reservoir and a variable mitochondrial complement determined by depot-specific thermogenic demand [13, 14]. Excess caloric intake enlarges these reservoirs, expanding cell size until lipid is mobilised.

Formerly viewed as a passive energy store, adipose tissue is now recognised as an endocrine organ and a permissive niche for secondary ovarian-cancer deposits. Adipocyte size correlates positively with secretion of leptin and interleukin-6 (IL-6), a pro-inflammatory cytokine that promotes tumour proliferation, angiogenesis and metastasis [15, 16, 17, 18, 19]. Conversely, obesity suppresses adiponectin, a hormone whose low levels are associated with aggressive disease [20, 21]. Enlarged adipocytes also become hypoxic, leading to necrosis, macrophage infiltration and chronic inflammation [22].

These interactions between stromal adipocytes and epithelial tumour cells, have intensified interest in adipose tissue as a dynamic component of the tumour microenvironment, with signalling functions now recognised as critical drivers of cancer initiation, metastasis, and therapy resistance [23]. Among these signals, adipocyte-derived factors, particularly exosomes, serve as key mediators of crosstalk; Williams et al. showed that exosomes from both malignant and benign adipocytes carry oncogenic signals that foster epithelial–to-mesenchymal transition (EMT), invasion, and chemoresistance in ovarian cancer cells [24]. Coupled with rising obesity rates, these insights add further complexity to adipose–tumour interactions and underscore the need to unravel the fat–cancer axis and to target adipose metabolic and signalling pathways in cancer therapy [23].

*In-silico* mathematical models are an excellent way to artificially recreate *in-vitro* biological experiments while saving on costs, time, and resources. They can readily be modified between simulations to see the impact of changing certain parameter values or inhibiting various mechanisms. Currently, there is an abundance of ordinary/partial differential equation (ODE/PDE) models relating adipose tissue to cancer [25, 26, 27, 28, 29], of which two of the most widely accepted and cited are briefly described below.

Ku-Carrillo et al [25] created an ODE model linking immunity, obesity, and tumour size. The model initially includes three cell populations: immune cells, cancer cells, and normal cells. A later adaptation of this proceeds to include a fourth cell type: adipocyte cells. It incorporates the general dynamics of how each cell type population proliferates using logistic growth and affects the density of others. The paper proceeds to compare the equilibrium points of the tumour size and its heavy dependence on parameter values associated to the fat levels of the patient. Another model developed by Yildiz et al [26] uses fractional derivatives to represent the dynamics and includes tumour, normal, and fat cells. Further analysis of the model looks into the optimal control problem aiming to maximise the normal cells to tumour cells ratio, while also minimising the usage of the drugs over time due to the negative side effects associated with their application. The results found that chemotherapeutic drugs should be used at varying rates, again depending on the values of certain parameters linked with the patient’s obesity level.

ODE models are very good at approximating the populations of cell types as a whole throughout the domain over time. Since there is no spatial aspect in an ODE model, important features such as the distance between two cells or the amount of a substrate at a certain location in the microenvironment can not be tracked. Agent-based models are models in which each agent (the cell) is given a set of user-defined rules. For example, proliferation rates can be set to depend on oxygen concentrations, or apoptosis rate may depend on a drug concentration. They are excellent at capturing the heterogeneity of a tumour microenvironment, since each cell is treated independently and according to its own surroundings. Agent-based models can be easily adjusted by changing individual rules to allow more control and more detailed dynamics than ODE models. The spatial aspect is crucial to include due to the inter cellular dynamics, hence the need for agent-based models in this line of oncological research.

A previous model by Oliver et al [30] explored the role of the bystander effect on hypoxia driven epithelial-to-mesenchymal transition (EMT) in ovarian cancer tumours. This study builds upon these ideas, incorporating the bystander effect to capture the EMT and tumour growth dynamics with a specific focus on the role of adipose tissue. Seminal studies by Sutherland introduced the tumour spheroid model, revealing that these 3D structures replicate the spatial gradients of nutrients, oxygen, and metabolic by-products observed in solid tumour microenvironments [31]. We aim to use a 3D multicellular model to determine the adipocyte cell-tumour cell interactions and their response to chemotherapies. By studying the intracellular and inter-cellular interactions within a tissue, our multiscale mathematical model aims to help predict the expected outcomes in an *in-silico* setting.

To perform our *in-silico* simulations, various dynamics will be implemented into a Physicell modeling framework. The simulations will run on a 3D spatial domain that tracks the concentration of each substrate with user created initial and boundary conditions. The concentrations are recorded on a discrete mesh while the cells can move continuously across the domain. Different cell types can be placed and initialised, each secreting and uptaking different levels of substrates. The underlying dynamics of each cell’s volume, mechanics, and motility can be defined by the user and adapted to fit the model created. Data collected experimentally will be used to find the required parameter values which haven’t previously been reliably clinically estimated. Functional genomics will be incorporated by the properties of each cell being passed onto daughter cells.

## 3 Biological Experiment

*In-vitro* experiments were performed over four days using two cell lines [30]: OVCAR-3 and SKOV-3. 2D monolayers and 3D spheroid tumours were cultured and placed in the presence of either omental tissue-conditioned media (OT-CM) or an unconditioned media (UCM) control. The presence of OT-CM was found to increase cell proliferation in all scenarios other than in OVCAR-3 spheroids, in which unconditioned media showed the highest cell luminescence after 96 hours (See Figure 1).

**Figure 1.**
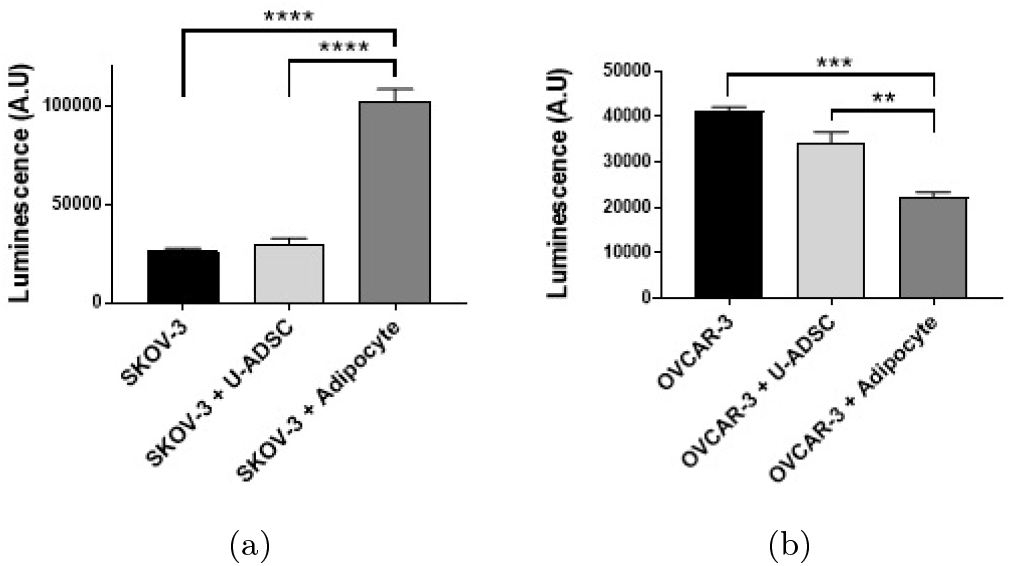
Final cell luminescence from the *in-vitro* experiments [30]. Cells are plated following co-culture with a control, undifferentiated adipose derived stem cells, and adipocytes. Populations of SKOV-3 (a) and OVCAR-3 (b) are taken after 96 hours, with the impacts of different types of media compared. SKOV-3 tumours co-cultured with adipocytes show an increased proliferation rate, while OVCAR-tumours co-cultured with adipocytes show a decreased proliferation rate.

The effects of undifferentiated adipose-derived stem cells (U-ADSCs) and adipocytes on tumour masses were also investigated. Further experiments tracked E-Cadherin, N-Cadherin, and Vimentin expressions throughout the tumour and microenvironment. It was found that the SKOV-3 tumour has a high band intensity of N-Cadherin and Vimentin, markers associated with a mesenchymal phenotype, suggesting epithelial to mesenchymal transition (EMT) had been completed (See Figure 2). The OVCAR-3 tumour showed a higher expression of E-Cadherin, thus suggesting that larger volumes of the neoplasm was made up of epithelial cells rather than mesenchymal. The cells on the periphery of the SKOV-3 tumour are shown to have E-Cadherin expression, suggesting an epithelial phenotype, while the interior cells appear to have a uniform, solely mesenchymal distribution. Mesenchymal OVCAR-3 cells are arranged in small clumps scattered within a surrounding pool of epithelial cells to form the solid tumour (See Figure 3).

**Figure 2.**
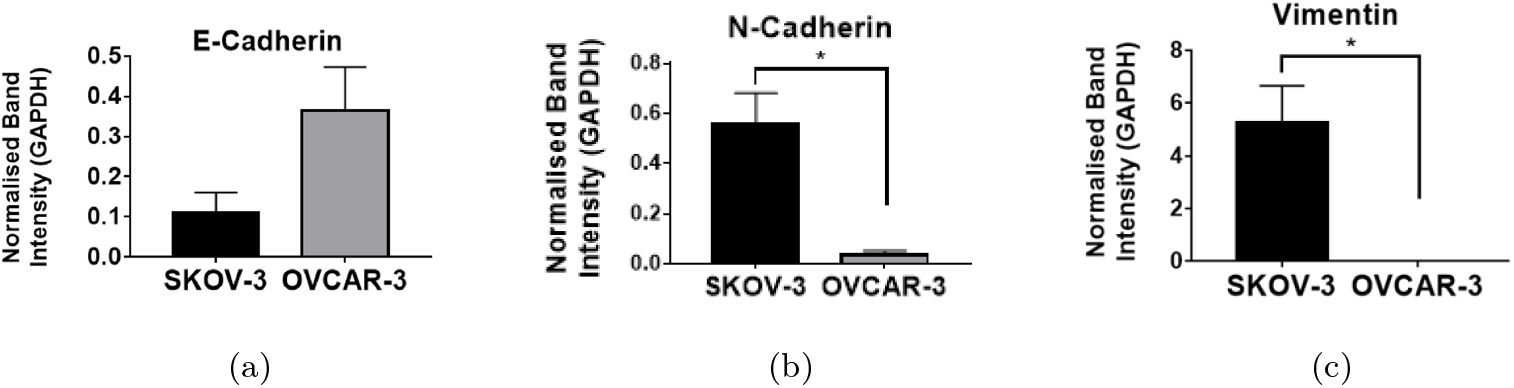
Normalised band intensity *in-vitro* between SKOV-3 and OVCAR-3 [30]. Epithelial markers (E-Cadherin) and mesenchymal markers (N-Cadherin and Vimentin) are expressed throughout the tumour after 96 hours. The intensities of these are normalised and compared for the different markets for SKOV-3 (black bars) and OVCAR-3 (grey bars) cell lines.

**Figure 3.**
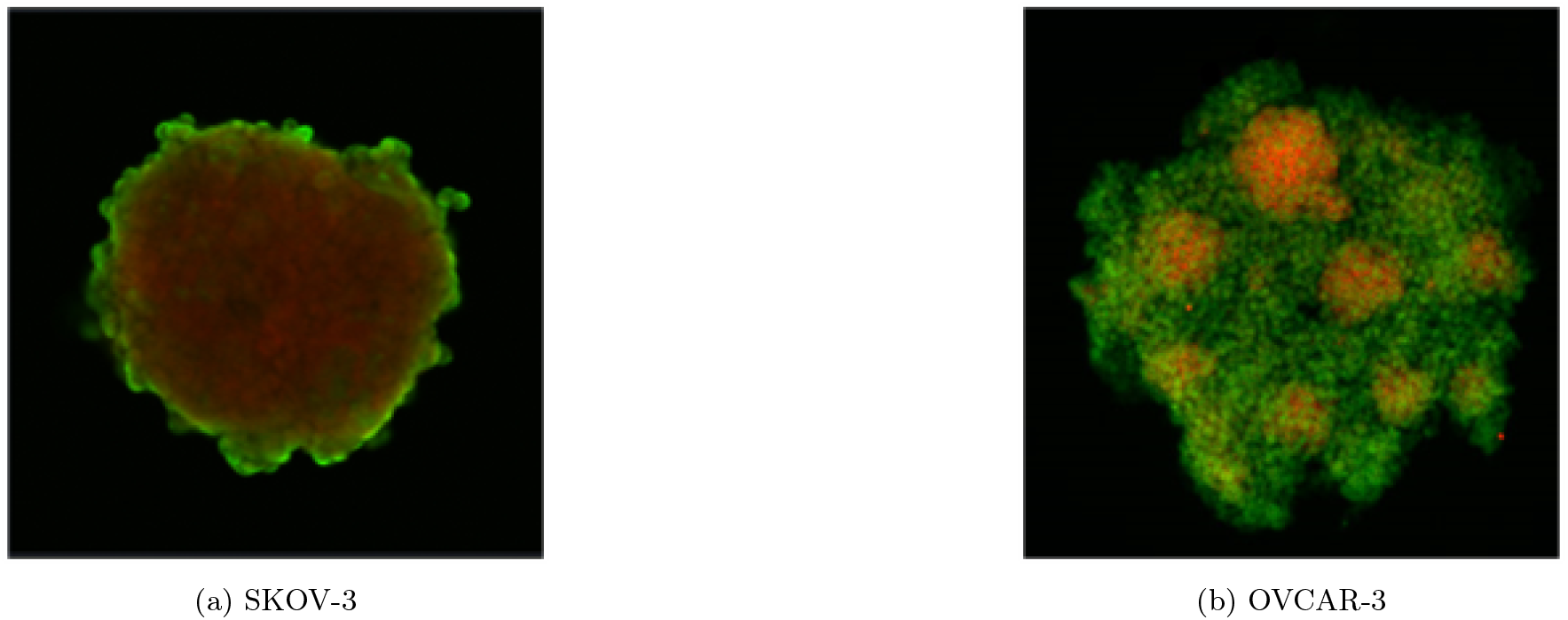
Final cadherin expressions in the cross-section of the tumours found *in-vitro* [30]. The spatial distribution of E-Cadherin (green) and N-Cadherin (red) is shown for SKOV-3 (a) and OVCAR-3 (b) spheroids after 96 hours.

The SKOV-3 spheroid in Figure 3 (a) shows a pool of red mesenchymal cells encased in a shell of green epithelial cells. This tumour has a consistent structure, with only mesenchymal cells expressing N-cadherin in the centre, and a thin layer of E-cadherin expressed by epithelial cells around the tumour periphery. In OVCAR-3 spheroids we observe a mass of primarily epithelial cells, with small clumps of congregated mesenchymal cells. Unlike SKOV-3 tumours, these mesenchymal cells are capable of forming through the tumour, including the periphery. These clumps of mesenchymal cells are thought to play a key role in metastasis, with the collective migration of the cells allowing these clumps to leave the primary tumour location and relocate elsewhere.

Figure 2 shows the expressions of E-cadherin (a), N-cadherin (b), and vimentin (c) in SKOV-3 and OVCAR-3 tumours. SKOV-3 spheroids show a low expression of E-cadherin, along with a high expression of N-cadherin and vimentin. This supports the image shown in Figure 3 (a), implying a mostly mesenchymal makeup of the SKOV-3 spheroid due to the tumour interior. Cells in OVCAR-3 tumours generally exhibit a high expression of E-cadherin and low expression of N-cadherin and vimentin, suggesting a largely epithelial population in OVCAR-3 tumours. This is in agreement with the cross sections of the tumour shown in Figure 3 (b), in which the majority of the tumour appears green due to expression of E-cadherin.

SKOV-3 cells appear to have a low proliferation rate, with an initial decrease in live cell populations and a slow increase during the final 48 hours of the *in-vitro* experiments (Figure 4 (a)). Omental tissue conditioned media is shown to increase the cycling rate of SKOV-3 cells, with populations of SKOV-3 cells in unconditioned media consistently lower throughout the experiments. OVCAR-3 cells show opposite trends, with unconditioned media generally leading to the highest proliferation rate in OVCAR-3 cells (Figure 4 (b)). Proliferation rates of OVCAR-3 cells are also considerably higher than those of SKOV-3, with around a tenfold increase in population sizes across four days.

**Figure 4.**
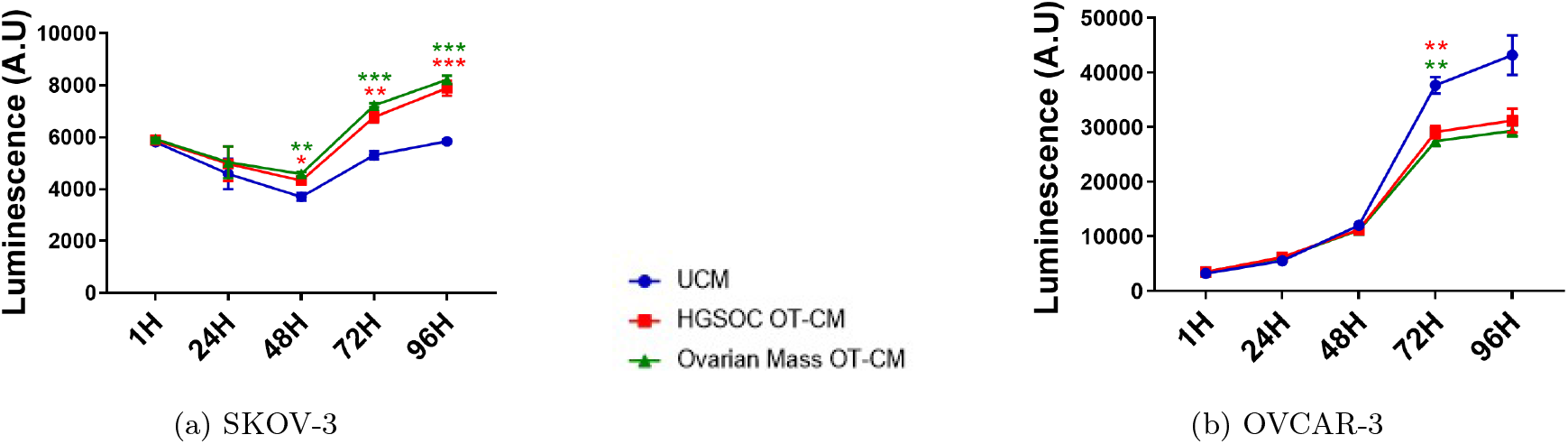
Temporal dynamics of SKOV-3 (a) and OVCAR-3 (b) tumours in various medias [30]. Live cell populations are measured daily for 96 hours in unconditioned media (blue), high-grade serous ovarian cancer omental tissue conditioned media (red), and ovarian mass omental tissue conditioned media (green). SKOV-3 tumours are initialised with 6000 cells and OVCAR tumours with 3000 cells.

The impact of adipocytes on the proliferation rate of SKOV-3 and OVCAR-3 cells is shown in Figure 1. Adipocytes increase the proliferation of SKOV-3 cells, causing a large increase in population size after 96 hours of the *in-vitro*. The presence of adipocytes has the opposite effect on OVCAR-3 tumours, in which unconditioned OVCAR-3 tumours show the highest final population while tumours plated with adipocytes show the lowest.

## 4 Model Framework

Here we develop an agent-based, multiscale model built upon a Physicell framework [32] to study this interplay between adipose tissue and cancer cells. Physicell is a physics based platform employed to generate biologically realistic simulations derived from user-defined rules and parameter values. There are two interacting layers to the simulation, one for the substrate concentrations and another for the cells (See Figure 5). BioFVM [33] is implemented to recreate the chemical microenvironment, in which multidimensional PDEs are implemented on a Cartesian mesh and solved using first order operator splitting (Equation (1)).

**Figure 5.**
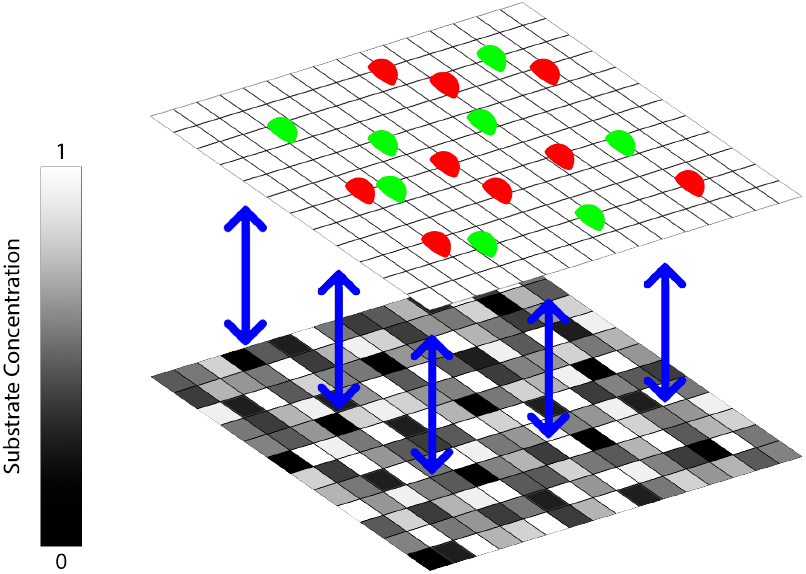
Example grid layout for a 2D simulation. Lighter shades on the bottom level of the visualisation show higher substrate concentrations diffusing according to Equation (1). Cells slide along a plane on the top level based on the value of the cell velocity, **v**_**i**_, given in Equation (2).

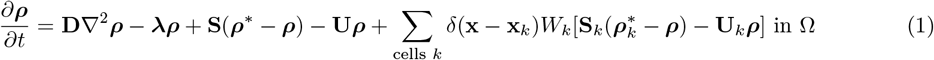

Here, ***ρ*** is the vector of chemical substrates, **D** and ***λ*** are the diffusion coefficients and decay rates respectively. **S** and **U** are the bulk supply and bulk uptake function respectively, while ***ρ***\* represents the vector of saturation densities for the substrates. The *δ*(**x**) term is the Dirac delta function, incorporated to include the source/sink points in the domain due to cell secretions/uptakes. *W*_*k*_, **x**_*k*_, **S**_*k*_, **U**_*k*_, and 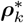 are the volume, position, source rates, uptake rates, and saturation densities respectively for the *k*^*th*^ cell.

The cells move off lattice within the 3D domain based on adhesion, repulsion, and various motility rules. The forces acting upon each cell are shown in Equation (2), used in the Physicell framework [32], giving the velocity after each time step and updating the position of cell *i* subsequent to each iteration using the second-order Adam’s Bashforth discretisation. Each cell has a repulsion force acting between itself and the surrounding cells to prevent overlapping. The speed of random motion is set to be two times higher for mesenchymal cells as it is for epithelial to qualitatively replicate the more metastatic behaviour of cells that have undergone EMT.

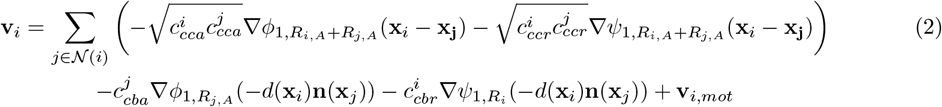

Here, 𝒩 (*i*) is the list of all cells in the domain capable of interacting with cell *i*. The magnitude of the cell-cell adhesion and repulsion forces are given by 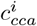 and 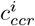 respectively, while the cell-basal membrane forces are given by 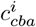 and 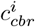. *R*_*i*_ is the radius of cell *i* and *R*_*i,A*_ its maximum adhesion distance. Functions *ϕ*_*n,R*_(**x**) and *ψ*_*n,R*_(**x**) are adhesion and repulsion functions, the details of which are defined in Physicell [32]. Finally, **n**(**x**) represents the normal vector to the nearest cell basal membrane and *d*(**x**) the distance to it.

### 4.1 Cell Cycle and Death

Cells in the model carry out four stages of the cell cycle before completing mitosis and dividing. The probability a cell progresses from stage *i* to *j* during an iteration is given by *≈ r*_*ij*_Δ*t*, where *r*_*ij*_ is the transition rate from state *i* to *j* and Δ*t* is the change in time. The presence of media is assumed to increase the proliferation of SKOV-3 cells by increasing the rate a cell leaves the G1 stage of the cell cycle (See Figure 6 (a)) [34]. The rate of proliferation of OVCAR-3 cell is decreased in the presence of media, as shown in Figure 6 (b). Mesenchymal cells are assumed to cycle 50% slower than epithelial cells, inferred from the biological experiments. SKOV-3 cells are also assumed to have a slower proliferation rate than OVCAR-3, spending approximately ten times as long in the G1 phase, thus increasing their total cycle rate approximately fivefold. The average time spent in each stage of the cell cycle is given in Tables 1.

**Table 1:**
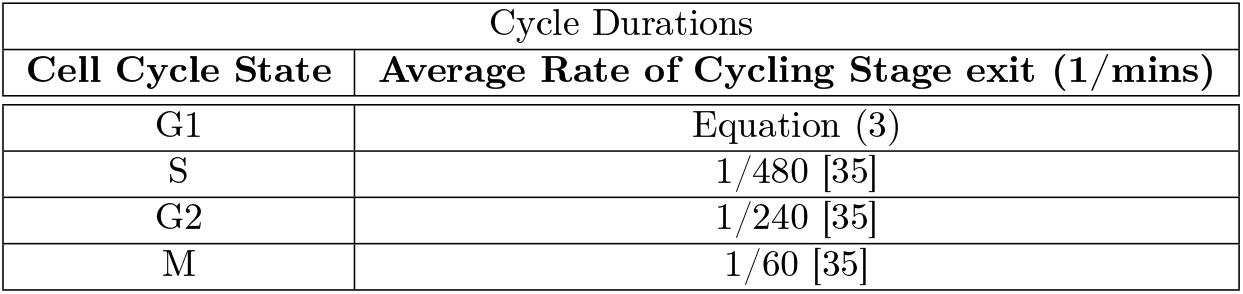
Average Cycle Durations.

**Figure 6.**
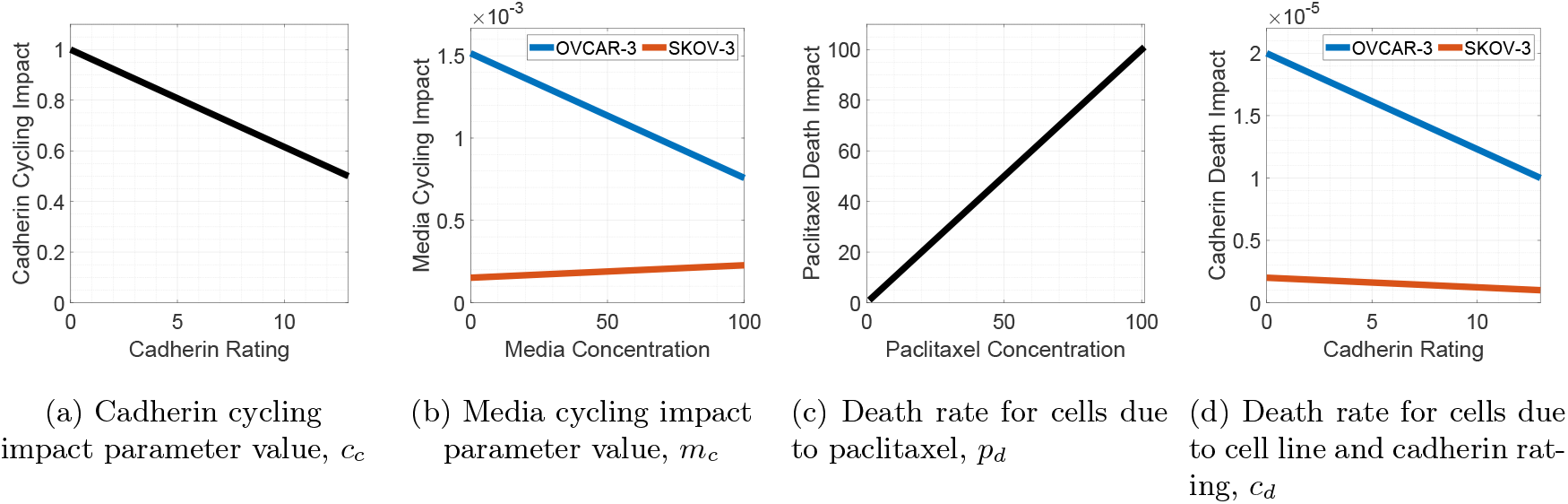
G1 cycling and death rates for OVCAR-3 and SKOV-3 cells. The relationships between cadherin rating and cycling rate (a), media concentration and cycling rate (b), paclitaxel concentration and apoptosis rate (c), and cadherin rating and apoptosis rate (d) are shown. OVCAR-3 relationships are shown in blue, SKOV-3 relationships are shown in red, and those independent of the cell lines are shown in black.

As slower cycling cells are assumed to reach cycling checkpoints less frequently than faster cycling cells, the death rate is also assumed to be lower. The lifespan of cells varies greatly from a timescale of days to years [36], so in our model we assume an OVCAR-3 epithelial cell will have an average lifespan of around 10 weeks. We assume that SKOV-3 cells cycle and die ten times as slowly as OVCAR-3 cells.

Figure 6 shows the rate for different cell phenotypes in varying media conditions to leave the G1 stage of the cell cycle and perform apoptosis. Cells with a more mesenchymal like phenotype cycle slower, incorporated into the model using Figure 6 (a). Based on an inspection of the experimental data shown in Figures 1 and 4, we assume for simplicity that a lack of media increases the proliferation rate of SKOV-3 cells [37] by 50% and decreases OVCAR-3 cells in spheroids [24] by 50%. This difference allows us to capture the phenomenon of adipose derived media increasing the tumour cell population compared to the control in SKOV-3 tumours, while decreasing it in OVCAR-3 tumours. These two impact parameters are incorporated into rate at which a cell leaves the G1 stage of the cell cycle using Equation (3).

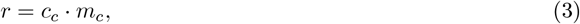

We will later compare the tumour growth with and without chemotherapy treatment. When treatment is included, paclitaxel is added uniformly throughout the domain after 24 hours, and all voxels are assigned a paclitaxel concentration of 50 (dimensionless). Paclitaxel is assumed to remain at a constant concentration throughout the course of the experiment, with higher concentrations in the microenvironment leading to higher rates of apoptosis in the cells, as shown in Figure 6 (c) [38]. Treatment in the model is assumed to lead up to a 100-fold increase in apoptosis rate. Epithelial cells are more susceptible to treatment due to their faster cycling rate, while the mesenchymal cells are more resistant [39]. We therefore assume for the model that epithelial cells have an apoptosis rate twice as high as mesenchymal cells (See Figure 6 (d)). These two impact parameters are incorporated into rate at which a cell can undergo apoptosis using Equation (4).

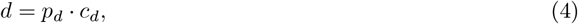

### 4.2 Role of the Bystander Effect

Several studies have investigated the cellular composition of patient-derived ovarian cancer spheroids. Capellero et al [40] reported that ascites-derived spheroids are composed predominantly of ovarian cancer cells, with minimal fibroblast or leukocyte contamination. Notably, these tumour cells exhibit a hybrid EMT phenotype, co-expressing epithelial markers such as EpCAM and cytokeratins alongside mesenchymal markers like *α*SMA and fibronectin [40]. This partial EMT state is thought to enhance spheroid viability and adhesion within the peritoneal cavity. EMT not only promotes cancer cell motility and dissemination by disrupting intercellular junctions but also confers stem-like traits that aid survival [41], while the reverse process, mesenchymal-to-epithelial transition (MET), is essential for metastatic re-differentiation and colonisation [42].

Building on these findings, we developed a model that incorporates the emergence of small mesenchymal clusters within a predominantly epithelial population of OVCAR-3 cells, as these clusters are crucial for faithfully replicating spheroid behaviour. In our model, we simulate a bystander effect, whereby epithelial cells near mesenchymal cells upregulate cadherin expression in response to local signalling cues, reflecting the plasticity of EMT/MET transitions and the influence of intermediate cell states on surrounding epithelial populations [43, 44]. The bystander effect has been observed in various biological processes, such as radiation induced damage [45, 46] or cell senescence being passed onto neighbouring cells [47]. To achieve this phenomenon, we include another substrate into the model. Mesenchymal cells which have undergone EMT are set to secrete a bystander signal [48] up to a rate of 100 units per minute, representing the percentage of the maximum secretion possible by cells (Figure 7).

**Figure 7.**
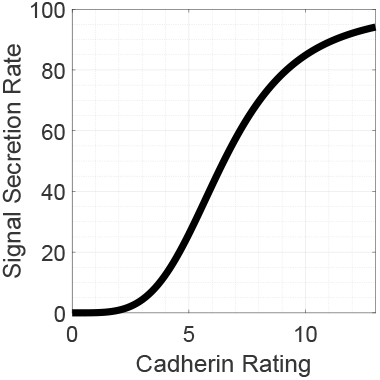
Signal secretion rate for cells dependant on their current cadherin rating. Cells at a higher cadherin rating with a more mesenchymal-like phenotype encourage epithelial cells in their proximity to undergo EMT. This is done by secreting a signal with low diffusion into the microenvironment to increase the rate of EMT.

The voxel in which a mesenchymal cell is located almost instantaneously fills with this signal due to the high secretion rate and remains so as long as the mesenchymal cell resides within it. We have a low substrate diffusion coefficient (1 micron^2^/min) for the signal to keep the range of influence low and include a decay rate for the signal of 1.0 units per minute to avoid a build up of signal in voxels which no longer host mesenchymal cells. A hill function is used to ensure there is a small secretion rate for cells of a low cadherin rating while still allowing a high secretion rate for mesenchymal cells. This hill function has a half max set at 6.5, since this is half the value of the maximum cadherin rating (13) cells can reach during the simulation. The saturation value is set to 100 units, at which point signal saturation is assumed to be reached. The hill power is set to four, optimised across simulations using observations from the experimental images shown in Figure 3.

Figure 8 shows an example of four time steps of mesenchymal (red) and epithelial (green) cells in a 10×10 grid. Each square in the grid represents a voxel in the domain. Cells move freely and continuously across the lattice. The arrows from the cell in the diagram show where the cell is moving and the compartment of the grid that the cell will be in at the next time step. The shade of a voxel shows the concentration of the signal at the time point (black indicates maximum signal and white indicates no signal). When an epithelial cell enters a square with a high signal concentration, the probability it transforms to mesenchymal is greatly increased (see Equation (5)). When the grid square is vacant of mesenchymal cells the signal immediately begins to decay from that compartment and the concentration decreases until it is below the threshold signal value, meaning the signal is no longer sufficient to increase a cell’s ability to perform EMT in that voxel.

**Figure 8.**
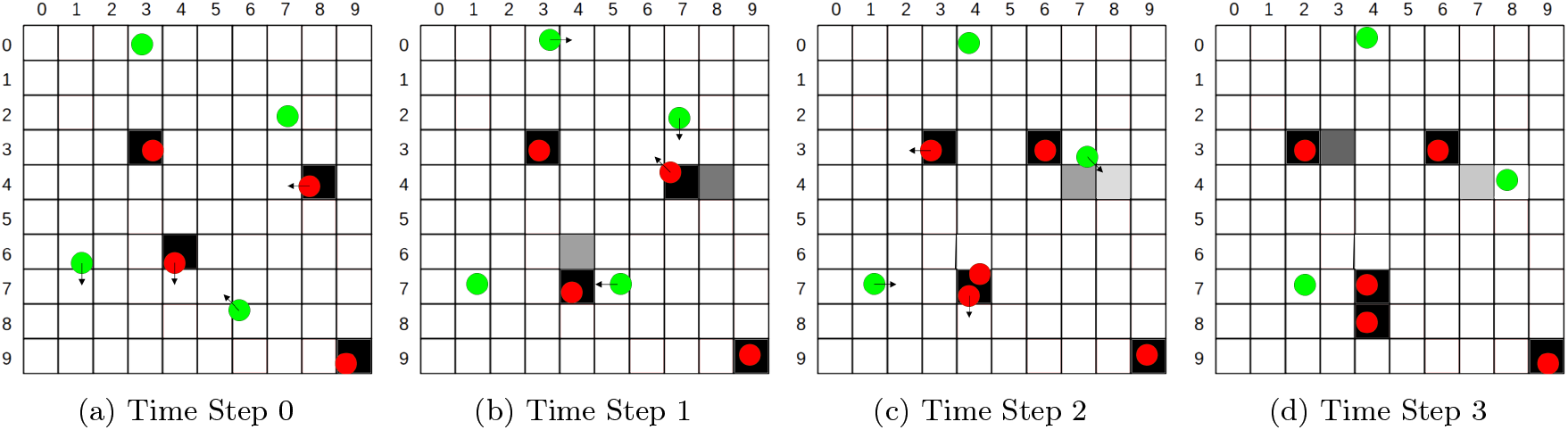
Example of the signal concentration during a simulation. Red mesenchymal cells secrete a chemical signal, shown by darker shades in the voxel. This signal encourages EMT in epithelial cells according to Equation (5). The signal slowly decays over time following the departure of the mesenchymal cell from the voxel.

Any cell with a rating of 7 or higher is considered to have completed EMT and adopts the behaviour assigned to a mesenchymal cell. We ensure the cells on the periphery of the SKOV-3 3D spheroids remain epithelial by incorporating a “transformation threshold” parameter, set in the model to be 0.1mmHg. Any SKOV-3 cell in oxygen levels greater than this value is assigned a cadherin rating of 0. This prohibits the cells on the exterior of the tumour from undergoing EMT and ensures they remain epithelial to replicate the biological experiments in which a shell of epithelial cells is wrapped around an interior mass of mesenchymal cells.

### 4.3 Intracellular and Intercellular impacts on EMT

Cells show the highest phenotypic plasticity when only partial EMT has been completed, with the most stable cells found to be either purely epithelial or purely mesenchymal. To include this, a cadherin EMT impact parameter, *c*_*e*_, is introduced (Figure 9). Along side this, cells in low levels of oxygen are more likely to jump up a cadherin rating and perform EMT than those which are not [49]. We introduce an oxygen EMT impact parameter, *o*_*e*_, in Figure 9 (b). The oxygen EMT impact parameter decreases as oxygen levels increase, saturating at zero when oxygen concentration increases above 1mmHg. Adipose derived media and signal secreted by mesenchymal cells are both assumed to encourage EMT in epithelial cells. This phenomenon is incorporated into the model using a media EMT impact parameter, *m*_*e*_, and a signal EMT impact parameter, *s*_*e*_. These parameters are positively correlated to the concentration of their respective substrate, with saturation being reached at concentrations of 100 units of media and 20 units of signal.

**Figure 9.**
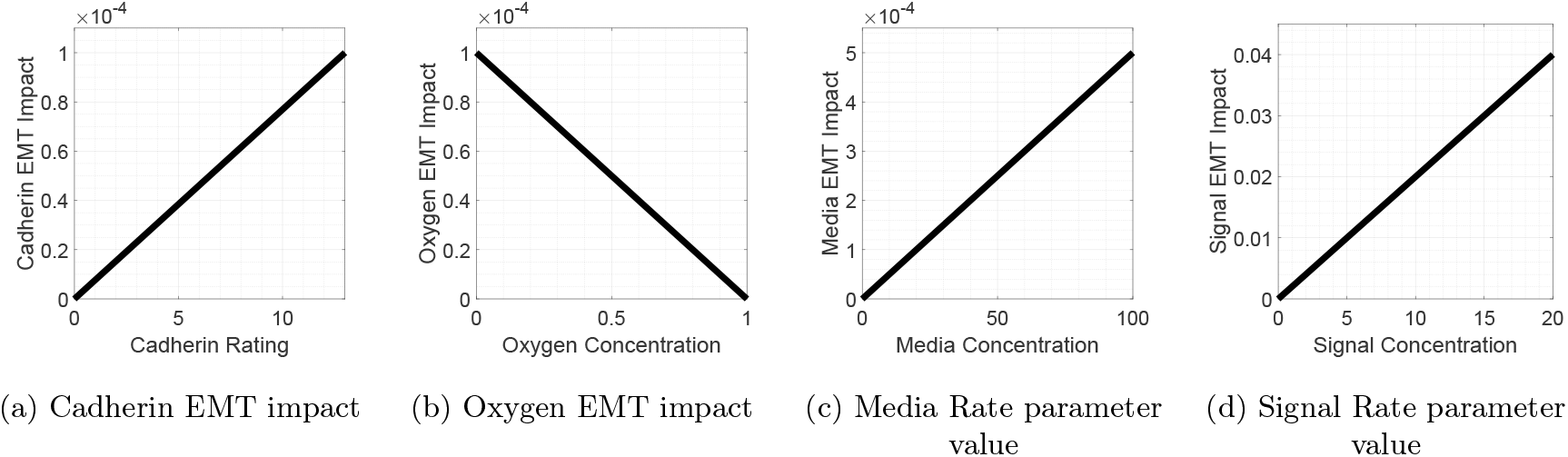
EMT rate variables with respect to the intra and inter-cellular conditions. The impact of the cadherin rating (a), oxygen concentration (b), media concentration (c), and signal concentration (d) on the rates of EMT IN cells. These variables are included in calculating the probability of EMT on each iteration using Equation (5).

All impact parameters are incorporated into calculating the probability for a cell to jump between cadherin ratings, thus ensuring the conditions lead to an impact on the rates of EMT. The parameter values are added together for each cell and this sum produces a jump probability shown in Equation (5). Here, *p* is the probability that a cell will jump up a cadherin rating on each six minute iteration.

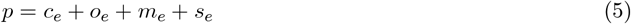

### 4.4 Parameter Values

The key parameters for our model, such as cell-cell adhesion strengths and substrate secretion/uptakes are stated in Tables 2 and 3. For simplicity, we keep cell volumes, repulsion and adhesion strengths, and substrate uptake rates equal between epithelial and mesenchymal cells. Cell adhesion (0.4*µm*/min) and repulsion (10.0*µm*/min) strength parameter values are kept consistent with those used in the Physicell template project [32]. These values help ensure there is no overlapping of the cell membranes whilst also preventing the tumour from excessively spreading out and not remaining as one collective mass.

**Table 2:** Cell Parameter Values.

| Parameter | Units | Value |
| --- | --- | --- |
| Cell Volume | microns<sup>3</sup> | 3000 [50] |
| Cell Adhesion Strength | micron/min | 0.4 |
| Cell Repulsion Strength | micron/min | 10 |
| Motility Speed |  | 0.1 [51] |
| Oxygen Uptake rate | mmHg/min | 0.6 [52] |
| Media Uptake rate | 1/min | 0.01 |

**Table 3:**
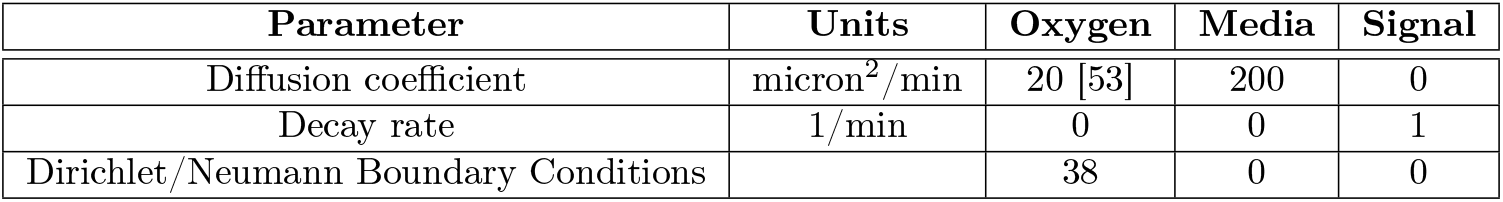
Substrate Parameter Values. Boundary conditions which are non zero use Dirichlet boundary conditions and those which are zero use Neumann boundary conditions with zero flux.

| Parameter | Units | Oxygen | Media | Signal |
| --- | --- | --- | --- | --- |
| Diffusion coefficient | micron<sup>2</sup>/min | 20 [53] | 200 | 0 |
| Decay rate | 1/min | 0 | 0 | 1 |
| Dirichlet/Neumann Boundary Conditions |  | 38 | 0 | 0 |

In simulations with adipocyte derived media, we initialise the domain with 50 dimensionless units of media distributed uniformly. Oxygen enters the system immediately using Dirichlet boundary conditions set to 38mmHg from each boundary edge. Epithelial and mesenchymal cells in both cell lines each uptake oxygen at a rate of 0.6mmHg per minute and media at a rate of 0.001 units per minute in the model, removing the respective amount of substrate from the cells voxel as a sink. Oxygen and media have diffusion coefficients of 20 micron^2^ and 200 micron^2^ per minute respectively. Due to the short time frame of the experiments we have assumed the decay rate of oxygen and media to be zero.

Cells have an oxygen uptake of 0.6mmHg/min in the *in-silico* model [52]. Despite the constant influx of oxygen incorporated from the boundary edges through Dirichlet boundary conditions, oxygen intake from the cells can create hypoxic conditions in the tumour core, resulting in a higher probability of EMT occuring. The adipose derived media is assumed to have a much smaller uptake rate than that of oxygen. The media has less direct involvement than oxygen for cellular respiration and so the cell uptake rate is set to be lower. We see in the biological experiments that the impact of media on the proliferation rate remains relatively high throughout the 96 hour *in-vitro* experiment, suggesting a low uptake rate of media in the cells. This 0.01 units per minute uptake rate ensures that central areas of the tumour can become media deficient and cells on the periphery proliferate the fastest as a result. The signal secretion rate is relatively redundant in this model as the threshold value for the signal to impact EMT rates is low. Any value larger than this threshold is sufficient to use and the changes in this secretion rate only change how long a voxel retains signal after a mesenchymal cell has departed it (See Table 2).

Oxygen and adipose derived media are assumed to each have no decay rate due to the short time frame of the simulation. The only reduction in the levels of these substrates is due to the uptake rate of the cells shown in Table 2 for processes such as glycolysis in respiration. The media is placed in the domain uniformly at the beginning of the simulation to immediately encourage cell proliferation. No further media is added during the simulation, resulting in a gradual decrease in overall media levels. There is an input of oxygen from the boundary edges to recreate the environment found in the *in-vitro* experiment. This is implemented using Dirichlet Boundary conditions along all faces of the 3D cube shaped domain. Media and signal substrates are assigned Neumann boundary conditions with a flux of zero along the boundary edges. We set the diffusion coefficient to be 20 for the oxygen substrate [53] and 200 for media. The decay rate of 1.0 units per minute for the signal substrate ensures the signal does not remain indefinitely in a voxel absent of mesenchymal cells but is not large enough to almost instantaneously reduce to 0 after one departs. Values for these parameters can be found in Table 3.

Table 4 shows the trends in cell behaviour between the two cell lines used in the simulations. OVCAR-3 cells are assumed to have a lower probability of performing EMT than SKOV-3 cells. This is observed in Figure 2 where SKOV-3 cell lines have a much higher expression of the mesenchymal marker N-Cadherin than OVCAR-3 cell lines. This goes hand in hand with the fact that OVCAR-3 cells are more likely to undergo mesenchymal to epithelial transition (MET). Since SKOV-3 spheroids are made up of a larger proportion of mesenchymal cells than OVCAR-3 spheroids, we assume not only that epithelial cells in SKOV-3 spheroids are more likely to transform to mesenchymal cells, but the reverse process of MET is less likely in SKOV-3 cells.

**Table 4:** Cell Type Differences.

| OVCAR-3 | SKOV-3 |
| --- | --- |
| Lower Probability of EMT | Higher Probability of EMT |
| All cells can perform EMT | Only Interior cells can perform EMT |
| Faster cell cycling rates | Lower cell cycling rates |

It is observed in the biological experiments in Figure 3 that the epithelial cells in any location of an OVCAR-3 spheroid are capable of performing EMT. This is unlike SKOV-3 spheroids in which the outside cells remain epithelial despite interior cells being primarily mesenchymal, implemented by including the oxygen threshold applied to SKOV-3 cells. OVCAR-3 is given no such threshold so that all cells have the potential to complete EMT. A final key difference between the cell lines is the impact that media has on cell proliferation. The increased cell luminescence with media in SKOV-3 spheroids suggests that media increases proliferation rates sufficiently to account for the generally slower proliferation rate of mesenchymal cells. This is not the case in OVCAR-3 cells despite the lower proportion of mesenchymal cells in the tumour. Therefore, we set the large impact of media on proliferation rates in SKOV-3 cells and a low impact on OVCAR-3 cells (See Table 4). Parameters not stated in this section were kept as the default Physicell parameter values from the template project provided [32].

## 5 Model Calibration

To ensure our model gave similar qualitative results to the biological experiments, we ran computer simulations to compare the tumour growth in unconditioned media and omental tissue conditioned media. Tumours are initialised as closely to the *in-vitro* experiments as possible. The simulated tumours were placed in 3D spheroids in a cube-shaped domain with edges of length 410*µm*. OVCAR-3 cells were initialised with 3000 cells in a sphere of radius 120*µm*, while SKOV-3 cells were initialised with 6000 cells in a sphere of radius 150*µm*. We ran the simulations for 96 hours of simulated time, storing cellular and microenvironment data every 60 minutes.

*In-vitro* SKOV-3 tumours shown in Figure 10 (a) are shown to have a large mass of mesenchymal cells in which there is a high expression of N-Cadherin creating the central red patch. The exterior shell of green epithelial cells is formed around the periphery. *In-vitro* OVCAR-3 tumours shown in Figure 10 (b) are shown to have small clumps of mesenchymal cells in the mass of otherwise epithelial cells. Both these trends are generally recreated in the results from the model simulations for SKOV-3 and OVCAR-3 spheroids shown in Figures 10 (c) and (d) respectively.

**Figure 10.**
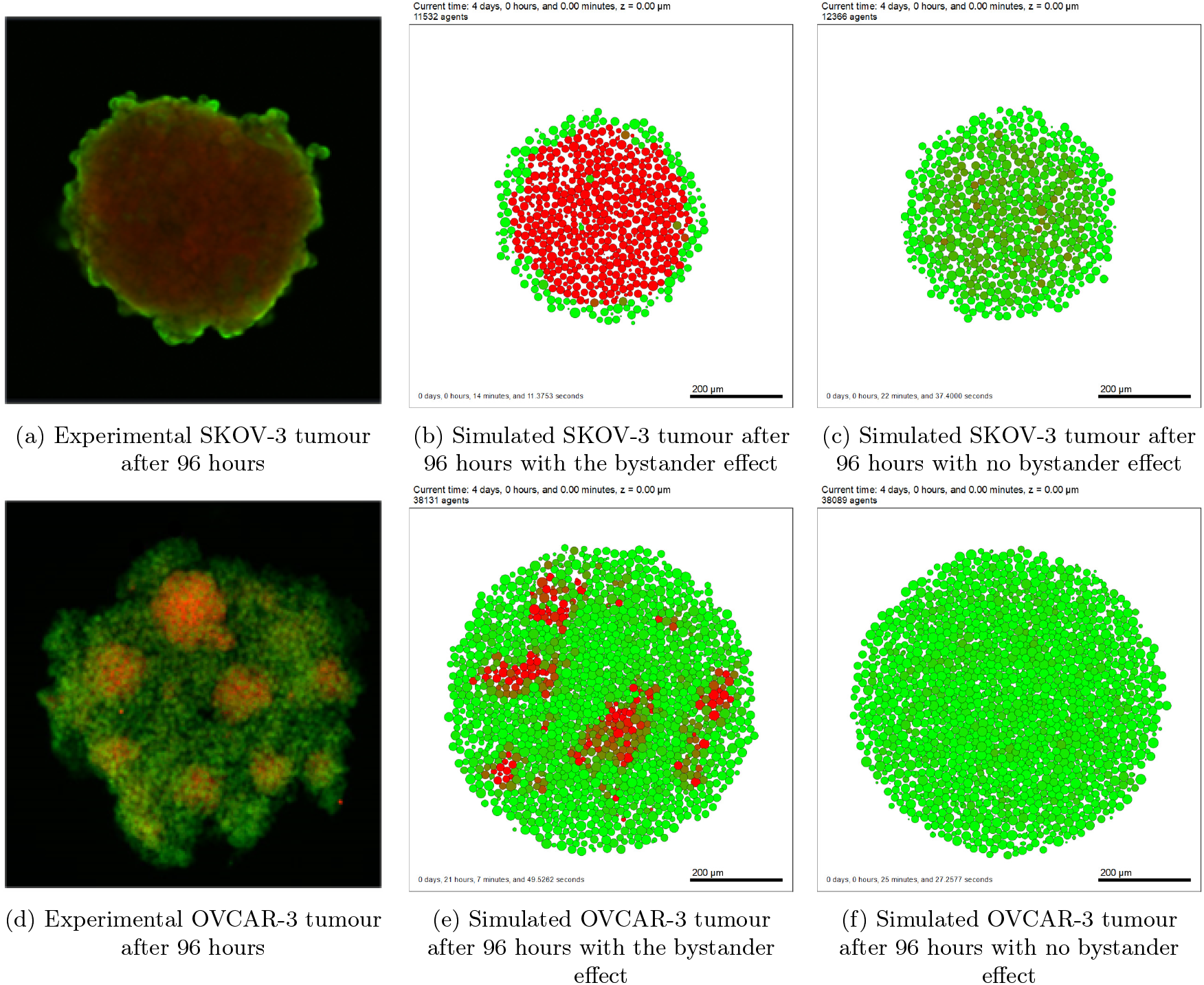
Comparisons of the final cadherin expressions in the cross-section of the tumours found *in-vitro* and *in-silico*. The spatial distribution of E-Cadherin (green) and N-Cadherin (red) is shown *in-vitro* for SKOV-3 (a) and OVCAR-3 (d) spheroids after 96 hours, and compared with those found *in-silico* for SKOV-3 (b) and OVCAR-3 (e). Simulations are also tested without the presence of the bystander effect in EMT, shown for SKOV-3 (c) and OVCAR-3 (f) tumours.

Figure 11 shows the live cell populations over time for the *in-silico* and *in-vitro* experiments in different types of media. The red line shows tumours grown in omental tissue-conditioned media, with the blue line representing tumour growth in unconditioned media. The green line shows tumour population grown in Ovarian Mass omental tissue-conditioned media, not used in the *in-silico* model for simplicity. Figure 11 (a) shows that omental tissue conditioned media increases cell proliferation compared with unconditioned media in SKOV-3 spheroids. Proliferation in OVCAR-3 spheroids is decreased when added to media in the *in-vitro* experiments, as shown in Figure 11 (b). The *in-silico* simulations for SKOV-3 and OVCAR-3 spheroids shown in Figures 11(c) and (d) respectively capture this general trend. While the exact shapes of the population curve during the simulation differ, the end result is generally similar between the biological experiments and model simulations.

**Figure 11.**
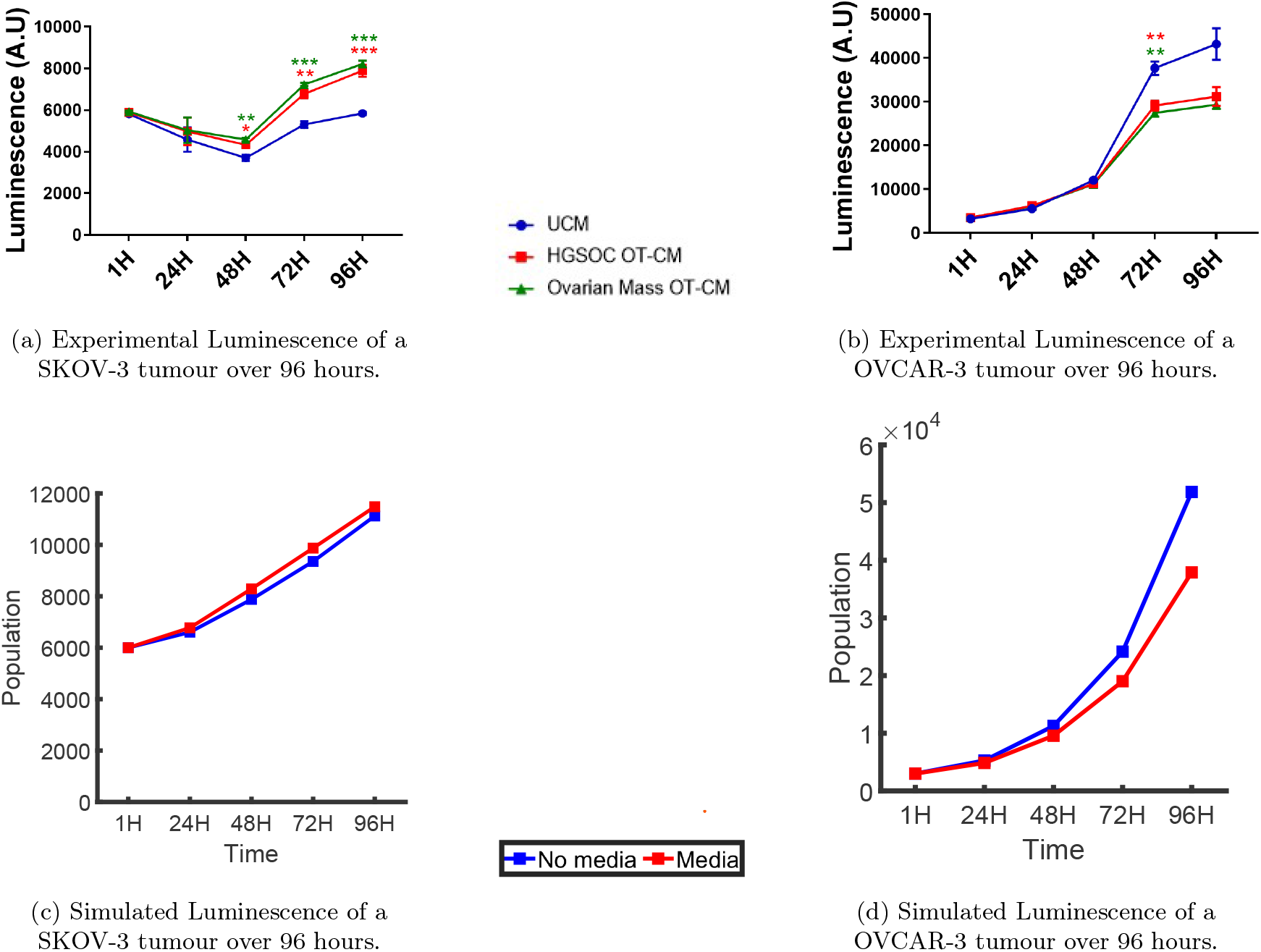
Comparison of temporal dynamics of tumours in various medias found *in-vitro* and *in-silico*. Results found from in vitro experiments are shown for SKOV-3 tumours (a) and OVCAR-3 tumours (b). Simulations are run for 96 hours in with no initial media concentration (blue) and with an initial media concentration of 50 units (red). These tumours are simulated for SKOV-3 cells (c) and OVCAR-3 cells (d), and compared with the results found *in-vitro* for the two cell lines.

We also compare the number of cells expressing mesenchymal and epithelial markers in the spheroids. After 96 hours of simulated time we calculated the proportion of mesenchymal cells and epithelial cells for OVCAR-3 and SKOV-3 tumours. The difference between the black bars in Figures 12 (a) and (b) suggest that SKOV-3 have a high N-Cadherin intensity and low E-Cadherin intensity, while OVCAR-3 tumours show very low N-Cadherin expression and high E-Cadherin expression following growth in adipose derived media.

**Figure 12.**
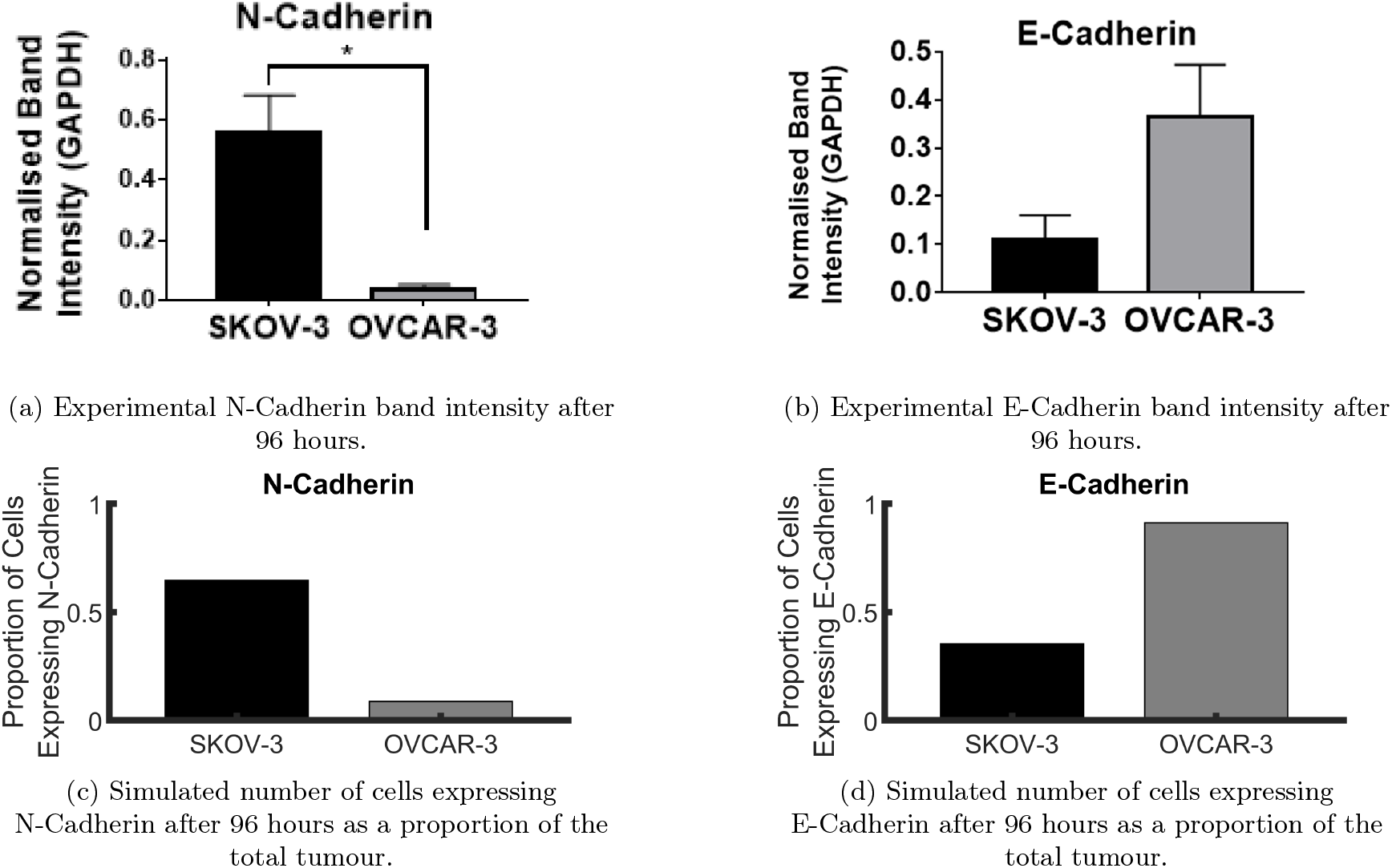
Comparison of the N-cadherin and E-cadherin expression found *in-vitro* and *in-silico*. Results found from *in-vitro* experiments show the expression of N-cadherin (a) and E-cadherin (b) after 96 hours. These are compared with the *in-silico* results for the two cell lines in which cells with a cadherin rated seven or higher are assumed to express N-cadherin, and cells with a cadherin rating lower than seven are assumed to express E-cadherin.

These results are also found in the computer simulations we perform. The results of these simulations shown in Figures 12 (c) and (d) fit well with the qualitative trends found in the *in-vitro* experiments.

This section helps to test the validity of the hypotheses we used to construct the model, with Figures 10, 11, and 12 showing similarities between the *in-silico* and *in-vitro* experiments. Figures 10 and 11 were used to set the assumptions of the model, with Figure 12 offering model validity using additional experimental data. General trends are consistently captured compared to the data provided from the *in-vitro* experiments. Ensuring our model has this calibration with the biological experiments provides the foundation to build from this model with the assumption that the results are accurate and realistic to what could be expected *in-vitro* or even *in-vivo*.

## 6 Results and Discussions

To explore the capabilities of the model, we begin to test the impact that various parameters can have on tumour growth and the final size of the neoplasm. We begin by comparing the effects of changing three different parameter values in the model. Firstly we vary the media in the initial conditions. This aims to show what may be expected of a tumour in patients varying from lean to obese. The previous sections have highlighted the importance of incorporating adipose tissue into a model and so looking into this impact in more detail beyond the *in-vitro* experiments could lead to a better understanding of the exact tumour attributes expected for different patients. We next introduce the treatment described in Section 4.1 to test the effectiveness of different drug dosages on a tumour. These results can be incorporated to study the dosage a patient should be administered, given their obesity level or composition of tumour. Finally, we compare the tumour growth for different initial sizes. This is a key determinant in what treatment should be given to a patient and stage diagnosis. This is especially the case when incorporating media since larger tumours will restrict the adipose derived media available to the central cells changing the dynamics over time. The details of these findings are highlighted below.

### 6.1 Effect of Media Concentration on Tumour Growth

To explore the impacts of adipose tissue and the omentum, we vary the levels of adipose derived media in the initial conditions of the microenvironment between different values. All parameters remain the same as stated previously other than the initial media concentration, assigned the default value of 50 dimensionless units distributed uniformly throughout the domain. We vary this parameter value between 0% and 200% of the default value shown in Figures 13 and 14, with the temporal dynamics of the populations compared.

**Figure 13.**
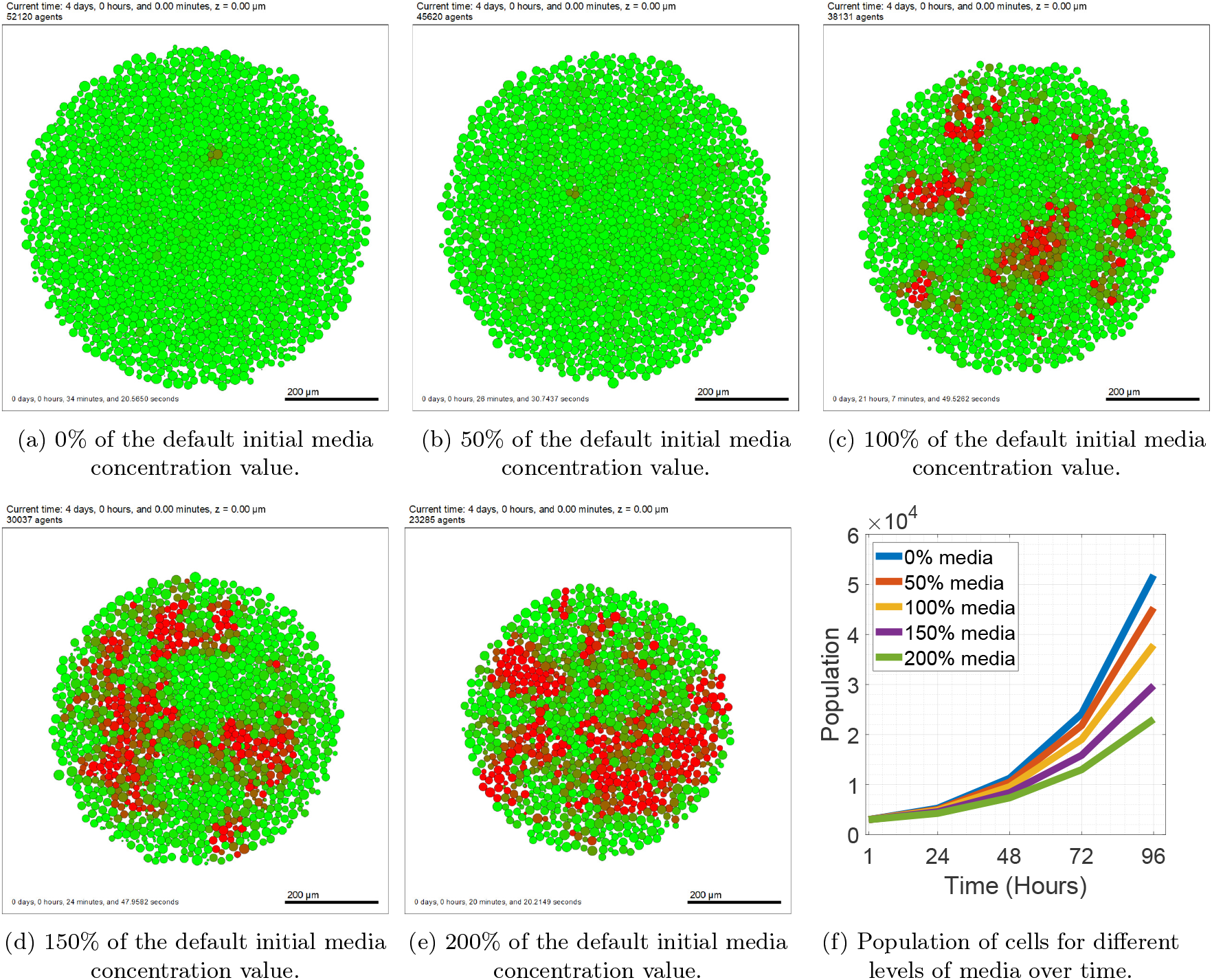
OVCAR-3 spheroids after 96 hours of simulated time for different levels of media concentration. A cross-section of the OVCAR-3 tumours after 96 hours are shown for spheroids initialised with no media (a), 50% of the default media concentration (b), 100% of the default media concentration (c), 150% of the default media concentration (d), and 200% of the default media concentration (e). The temporal dynamics of these simulations show the live cell populations over time (f).

**Figure 14.**
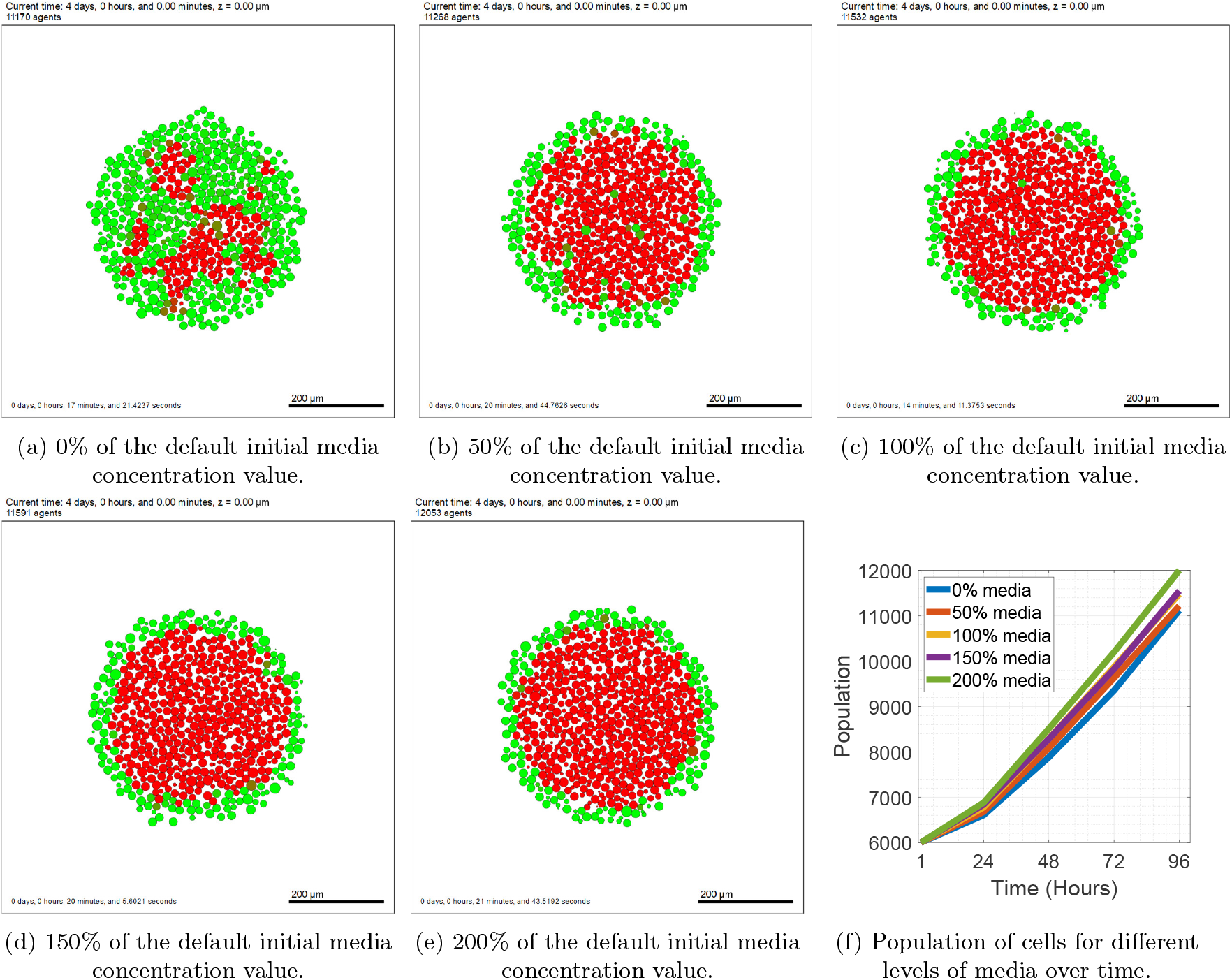
SKOV-3 spheroids after 96 hours of simulated time for different levels of media concentration. A cross-section of the SKOV-3 tumours after 96 hours are shown for spheroids initialised with no media (a), 50% of the default media concentration (b), 100% of the default media concentration (c), 150% of the default media concentration (d), and 200% of the default media concentration (e). The temporal dynamics of these simulations show the live cell populations over time (f).

#### 6.1.1 OVCAR-3

Figure 13 shows the tumour composition for OVCAR-3 spheroids after 96 hours in different media levels. For adipose derived media levels 50% of the default value and under ((a) and (b))there is insufficient media to encourage EMT to occur and the tumour ends with almost only epithelial cells present. The population of mesenchymal cells increases with levels of media and eventually the separate clumps begin to merge to form larger clumps as seen in Figure 13 (d) and (e). We see that the total population of cells at the end time of the simulation decreases monotonically as initial media conditions increase (f). While these results may suggest that increased fat levels may be beneficial due to less overall tumour growth over time, mesenchymal cells have a higher tendency to metastasise and are considered to be harder to control and treat than epithelial cells. Therefore, it is likely that tumours in lower adipose derived media are easier to treat despite their marginally larger size.

#### 6.1.2 SKOV-3

Figure 14 shows that only simulations with no adipose derived media remain fairly epithelial after 96 hours. All other initial conditions in the microenvironment lead to outcomes in which the interior of the tumour is almost entirely mesenchymal and the exterior is epithelial. Other than occasional individual cells, visually there are few major differences between (b), (c), (d), and (e) in Figure 14. Figure 14 (f) shows there is a small change in the total population of cells over time for different media conditions. Higher media consistently leads to a slightly larger tumour size at the end of the simulation.

### 6.2 Effect of Treatment Dosage on Tumour Growth

We next look for any potential impacts of administering treatment to a patient (Figures 15 and 16). We keep all parameters identical to those previously stated and introduce paclitaxel, highlighted in Section 4.1. We set the default full drug amount to be 50 units and vary this dosage between 0% (a), 50% (b), 100% (c), 150% (d), and 200% (e) of this default amount. The cell populations over time are also given in (f) for the different levels of drug dosage.

**Figure 15.**
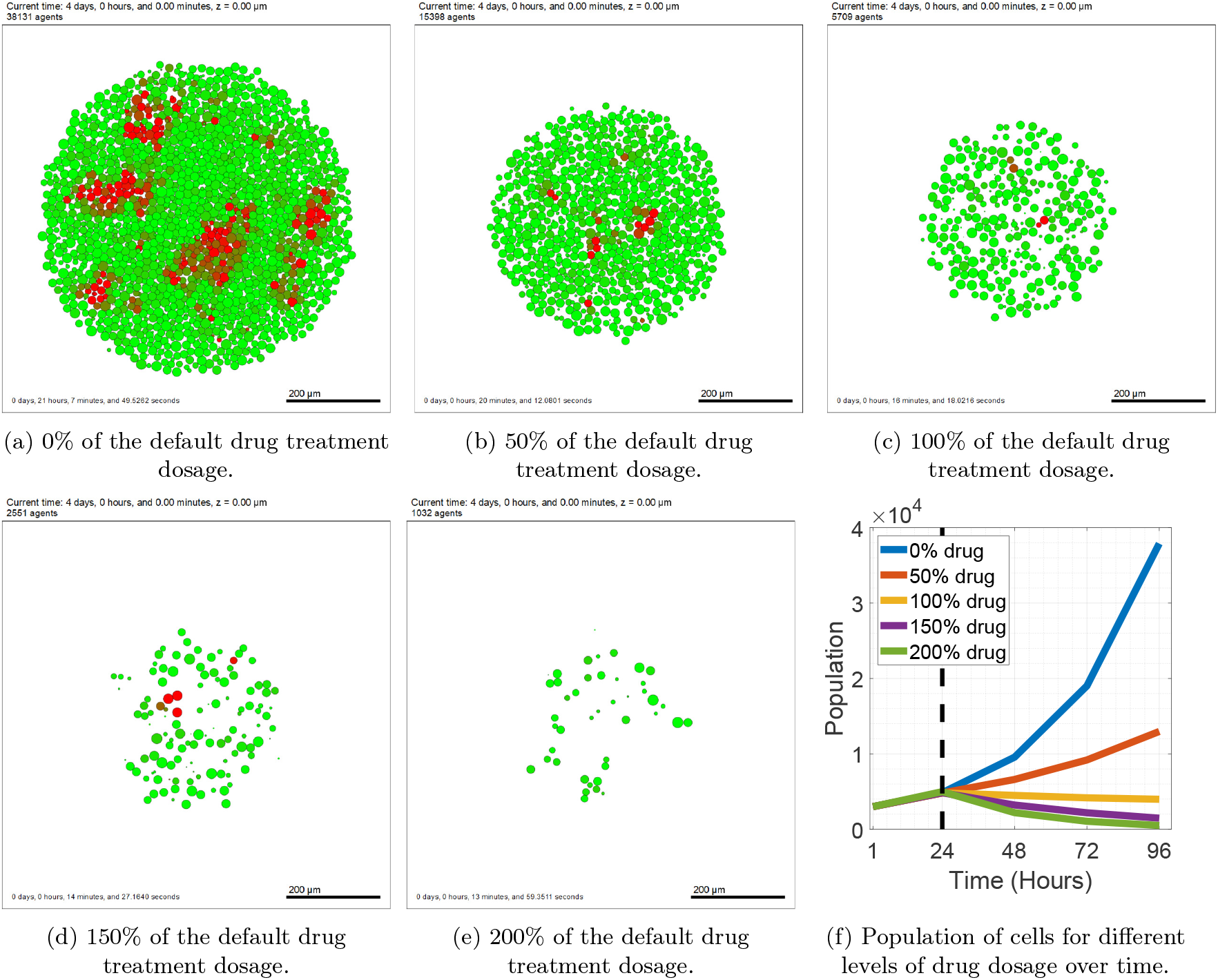
OVCAR-3 spheroids after 96 hours of simulated time for different levels of treatment. A cross-section of the OVCAR-3 tumours after 96 hours are shown for spheroids with no treatment (a), 50% of the default treatment dosage (b), 100% of the default treatment dosage (c), 150% of the default treatment dosage (d), and 200% of the default treatment dosage (e). The temporal dynamics of these simulations show the live cell populations over time, with the dashed vertical line showing the time of treatment introduction (f).

**Figure 16.**
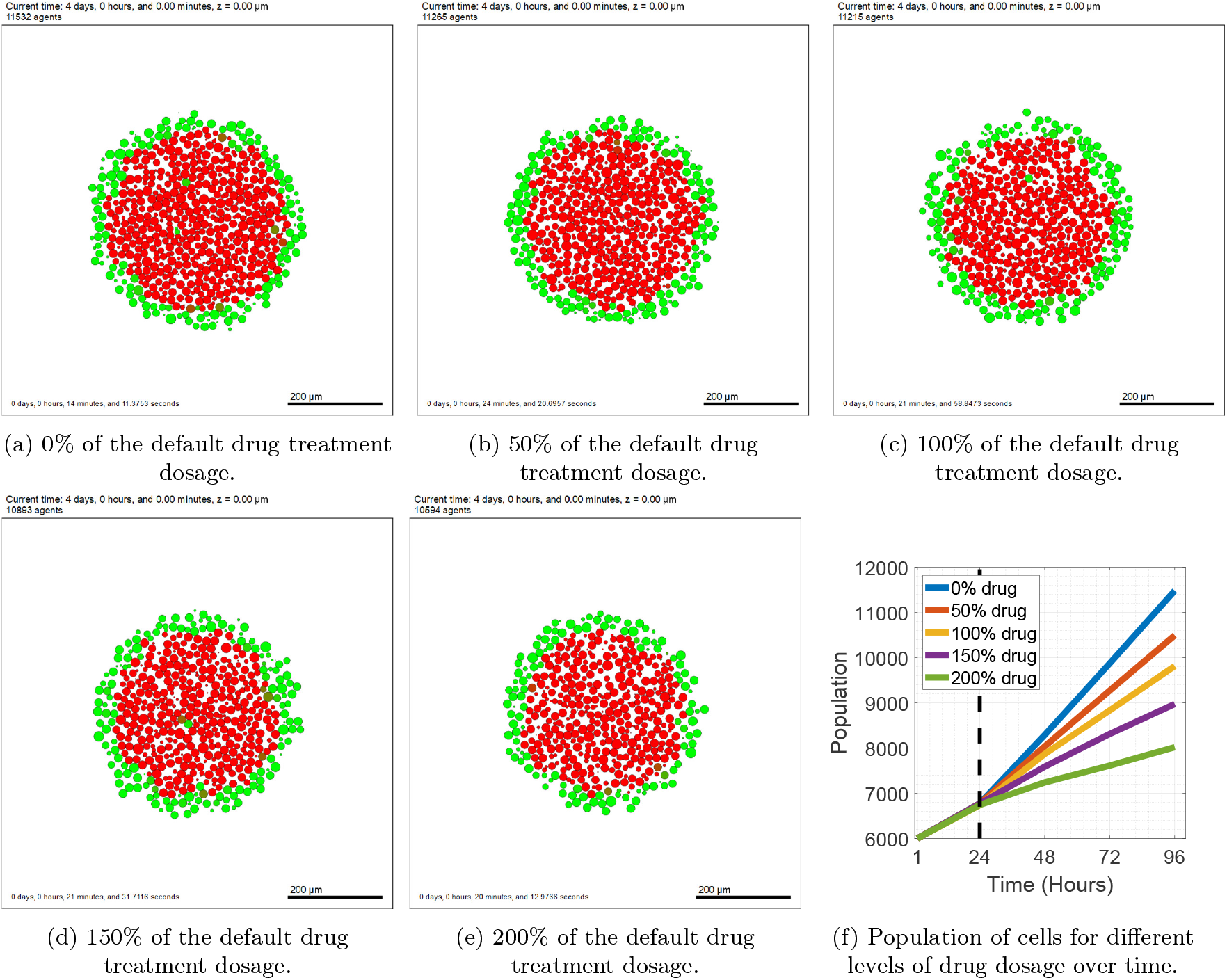
SKOV-3 spheroids after 96 hours of simulated time for different levels of treatment. A cross-section of the SKOV-3 tumours after 96 hours are shown for spheroids with no treatment (a), 50% of the default treatment dosage (b), 100% of the default treatment dosage (c), 150% of the default treatment dosage (d), and 200% of the default treatment dosage (e). The temporal dynamics of these simulations show the live cell populations over time, with the dashed vertical line showing the time of treatment introduction (f).

#### 6.2.1 OVCAR-3

Figure 15 shows the final OVCAR-3 spheroids after being exposed to different concentrations of paclitaxel during the simulations. No level of drug we tested was sufficient in eradicating the tumour entirely, despite leading to a significant decrease in size overall. The administration of paclitaxel has a lower impact on the death rate of the more resistant mesenchymal cells than on epithelial cells. This leads to the question of whether higher drug dosages are always better at killing a tumour, as it can leave more space open for the drug resistant, harder to treat cells to proliferate into. As a result, future treatments may be less effective in killing the remaining cells.

#### 6.2.2 SKOV-3

We next investigate the impact of the drug on SKOV-3 cells. Figure 16 shows the tumour appearance at the end of the simulations. Again, no treatment level tested in the simulations led to the complete removal of the tumour. The drug targets the epithelial cells on the periphery, stripping the tumour of its outside layer, thus increasing the oxygen levels to the previously interior cells. This converts the outside mesenchymal cells into epithelial cells through MET due to the increased oxygen levels available to these cells. As a result, the tumour shrinks over time while simultaneously keeping the lining of green epithelial cells on the tumour periphery. This process continues until the drug concentration is too low to increase the apoptosis rate of the epithelial cells, at which point the tumour can begin to recover.

No drug concentration tested for either cell line is sufficient to completely eradicate the tumour over time. While simulated tumour populations decrease with higher drug administration, certain drawbacks should be considered when treatment protocol is decided such as paclitaxel resistance, side effects of chemotherapy, and magnitude of the treatment effectiveness. In the future, we aim to test different drug types with different intervals between each administration to replicate current treatment protocols.

### 6.3 Effect of Initial Tumour Size on Tumour Growth

The next result we use our model for is to find is the importance of the initial tumour size on the cell population. Again, all parameters are kept as stated in previous sections other than initial number of cells. We have a default tumour size of 3000 cells for OVCAR-3 spheroids and 6000 cells for SKOV-3 spheroids (See Section 5). We also compare a small tumour size (a) (50% of this population), and a large tumour size (c) (150% of this population) and the populations over time across these three scenarios in (d). The difference in initial tumour size can provide an option for the model to account for the stage in which the tumour is diagnosed and how advanced the tumour is in the area of interest.

#### 6.3.1 OVCAR-3

Figure 17 shows the tumour growth for a small, medium, and large initial tumour size. Other than their size, the overall appearance of the tumours and the presence of the small mesenchymal clumps in the sphere of green epithelial cells remains similar. Since the larger tumours lead to a more restricted availability of media to each cell, the ratio of mesenchymal cells to epithelial cells decreases in larger OVCAR-3 spheroids. The rate of growth is similar between the three simulations and tumours starting twice as large in our model finish around twice as large after 96 hours of simulated time.

**Figure 17.**
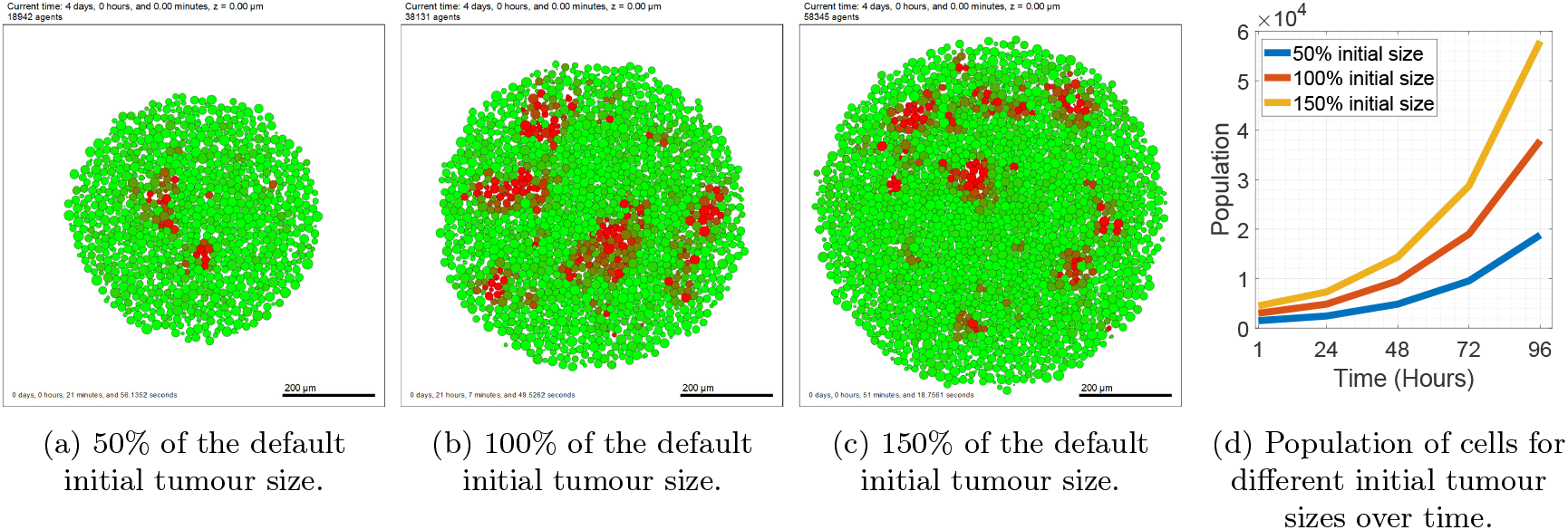
OVCAR-3 spheroids after 96 hours of simulated time for different initial tumour sizes. A cross-section of the OVCAR-3 tumours after 96 hours are shown for spheroids with 50% of the default initial population size (a), 100% of the default initial population size (b), and 150% of the default initial population size (c). The temporal dynamics of these simulations show the live cell populations over time (f).

#### 6.3.2 SKOV-3

Similar to OVCAR-3 spheroids, SKOV-3 spheroids shown in Figure 18 appear comparable throughout the variations of initial tumour size other than overall cell populations. For all different values of initial cell population we consistently have a thin layer of green epithelial cells on the tumour periphery and a red pool of mesenchymal cells in the interior after 96 hours of simulated time. The growth rates between the variations are all similar and can be accurately estimated to be used in treatments, given the initial size of the tumour.

**Figure 18.**
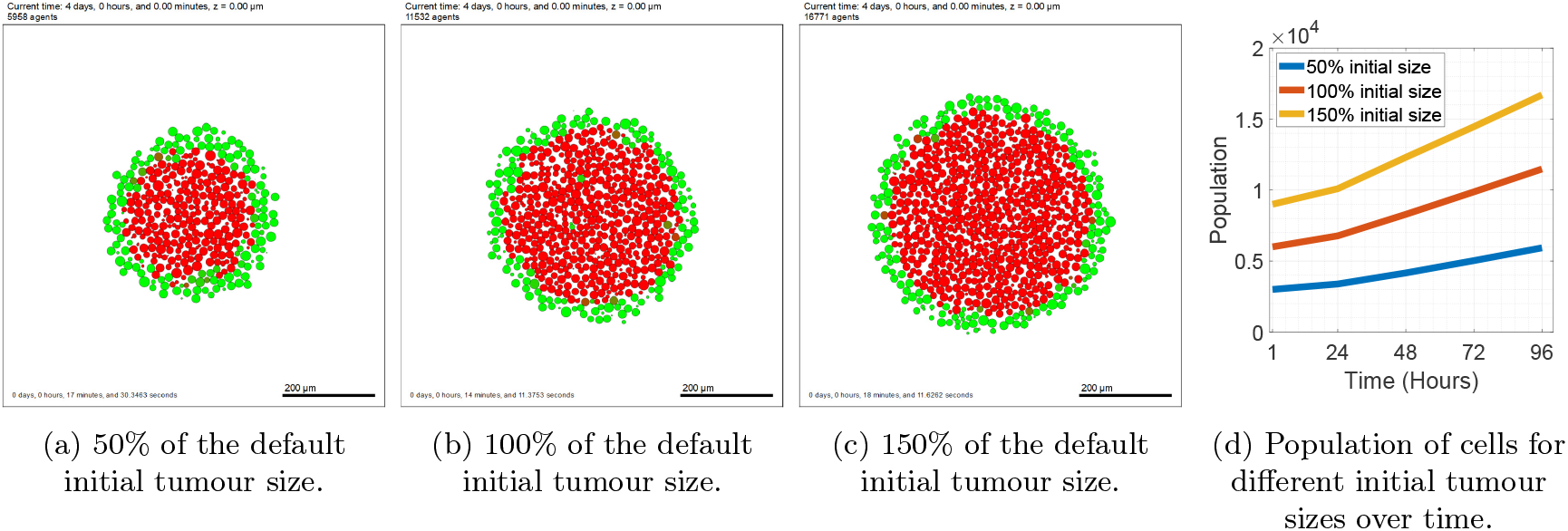
SKOV-3 spheroids after 96 hours of simulated time for different initial tumour sizes. A cross-section of the SKOV-3 tumours after 96 hours are shown for spheroids with 50% of the default initial population size (a), 100% of the default initial population size (b), and 150% of the default initial population size (c). The temporal dynamics of these simulations show the live cell populations over time (f).

## 7 Conclusions

The impact that adipose tissue has on cancer patients is clearly shown to be crucial, with fat tissue encouraging EMT and cell proliferation in both epithelial and mesenchymal cells. The exact mechanisms responsible for this are still largely unknown but can be represented in mathematical equations used for the simulations to ensure the model represents a cancer tumour as closely as possible. Section 5 showed that the model can recreate trends found in biological experiments with accurate results. This validation helps justify our ability to make predictions and gain answers to questions not yet found through *in-vivo* or *in-vitro* experiments. Unlike ODE models, the fact that the model is agent-based with a set of rules for each cell phenotype makes the model extremely adaptable, as a single rule can be modified based on biological justifications, without changing the dynamics of the entire system or other populations. The adaptability of this model allows for testing various plausible hypotheses that may play a role in EMT.

The flexibility of the model and capacity to generate custom tumours and microenvironments helps us take further steps towards studying tumour-specific treatment plans. In doing so, we hope to eventually be able to find the optimal treatment protocol for each patient, thus increasing the five year survival rate for ovarian cancer. In the future, the model can include additional agents such as fibroblasts and macrophages to ensure it is as close to *in-vivo* experiments as possible. Currently, the adipose derived media is considered to be a single collective substrate, which is not the case as it includes various types of peptides and exosomes [54]. When developing a mathematical model, it is important to strike a good balance between complexity and adaptability. For this reason, we create this model as a basis for future work and hope to explore increasing the complexity in future advancements and the pros/cons that come with doing so.

## Acknowledgments

SO was supported by EPSRC Maths DTP 2021/22 Swansea University [Grant EP/W523963/1]. MW was supported by Swansea University Texas academic partnership PhD programme.

## References

[1] G. C. Jayson, E. C. Kohn, H. C. Kitchener, and J. A. Ledermann, “Ovarian cancer,” The Lancet, vol. 384, no. 9951, pp. 1376–1388, 2014.

[2] M. H. Ebell, M. B. Culp, and T. J. Radke, “A systematic review of symptoms for the diagnosis of ovarian cancer,” American journal of preventive medicine, vol. 50, no. 3, pp. 384–394, 2016.

[3] L. Hardiman, “The case for early palliative care in the treatment of ovarian cancer,” Journal of the Advanced Practitioner in Oncology, vol. 5, no. 4, p. 290, 2014.

[4] W. Lee, S. Y. Ko, M. S. Mohamed, H. A. Kenny, E. Lengyel, and H. Naora, “Neutrophils facilitate ovarian cancer premetastatic niche formation in the omentum,” Journal of Experimental Medicine, vol. 216, no. 1, pp. 176–194, 2019.

[5] B. Topor, R. D. Acland, V. Kolodko, and S. Galandiuk, “Omental transposition for low pelvic anastomoses,” The American journal of surgery, vol. 182, no. 5, pp. 460–464, 2001.

[6] R. L. Siegel, A. N. Giaquinto, and A. Jemal, “Cancer statistics, 2024,” CA: a cancer journal for clinicians, vol. 74, no. 1, pp. 12–49, 2024.

[7] W. Gao, H. Yuan, S. Yin, R. Deng, and Z. Ji, “Identification of three subtypes of ovarian cancer and construction of prognostic models based on immune-related genes,” Journal of Ovarian Research, vol. 17, no. 1, p. 208, 2024.

[8] A. T. Ali, O. Al-Ani, and F. Al-Ani, “Epidemiology and risk factors for ovarian cancer,” Menopause Review/Przegl^1^d Menopauzalny, vol. 22, no. 2, pp. 93–104, 2023.

[9] C. M. Olsen, A. C. Green, D. C. Whiteman, S. Sadeghi, F. Kolahdooz, and P. M. Webb, “Obesity and the risk of epithelial ovarian cancer: a systematic review and meta-analysis,” European journal of cancer, vol. 43, no. 4, pp. 690–709, 2007.

[10] C. M. Olsen, C. M. Nagle, D. C. Whiteman, R. Ness, C. L. Pearce, M. C. Pike, M. A. Rossing, K. L. Terry, A. H. Wu, H. A. Risch, et al., “Obesity and risk of ovarian cancer subtypes: evidence from the ovarian cancer association consortium,” Endocrine-related cancer, vol. 20, no. 2, pp. 251–262, 2013.

[11] T. Wang, A. K. Sharma, and C. Wolfrum, “Novel insights into adipose tissue heterogeneity,” Reviews in Endocrine and Metabolic Disorders, pp. 1–8, 2022.

[12] W. Khazen, J.-P. M’Bika, C. Tomkiewicz, C. Benelli, C. Chany, A. Achour, and C. Forest, “Expression of macrophage-selective markers in human and rodent adipocytes,” FEBS letters, vol. 579, no. 25, pp. 5631–5634, 2005.

[13] S. C. Sarode, G. S. Sarode, N. Sengupta, N. K. Sharma, and S. Patil, “Adipocyte-tumor cell native encounter in oral squamous cell carcinoma,” 2020.

[14] M. Klingenspor, “Cold-induced recruitment of brown adipose tissue thermogenesis,” Experimental physiology, vol. 88, no. 1, pp. 141–148, 2003.

[15] S. Galic, J. S. Oakhill, and G. R. Steinberg, “Adipose tissue as an endocrine organ,” Molecular and cellular endocrinology, vol. 316, no. 2, pp. 129–139, 2010.

[16] F. Lönnqvist, L. Nordfors, M. Jansson, A. Thörne, M. Schalling, P. Arner, et al., “Leptin secretion from adipose tissue in women. relationship to plasma levels and gene expression.,” The Journal of clinical investigation, vol. 99, no. 10, pp. 2398–2404, 1997.

[17] M. Walter, S. Liang, S. Ghosh, P. Hornsby, and R. Li, “Interleukin 6 secreted from adipose stromal cells promotes migration and invasion of breast cancer cells,” Oncogene, vol. 28, no. 30, pp. 2745–2755, 2009.

[18] E. Dos Santos, F. Duval, F. Vialard, and M.-N. Dieudonné, “The roles of leptin and adiponectin at the fetal-maternal interface in humans,” Hormone molecular biology and clinical investigation, vol. 24, no. 1, pp. 47–63, 2015.

[19] Y. Shimada, Y. Komatsu, I. Ikezawa-Suzuki, H. Tai, N. Sugita, and H. Yoshie, “The effect of periodontal treatment on serum leptin, interleukin-6, and c-reactive protein,” Journal of periodontology, vol. 81, no. 8, pp. 1118–1123, 2010.

[20] T. Jardé, F. Caldefie-Chézet, N. Goncalves-Mendes, F. Mishellany, C. Buechler, F. Penault-Llorca, and M.-P. Vasson, “Involvement of adiponectin and leptin in breast cancer: clinical and in vitro studies,” Endocrine-related cancer, vol. 16, no. 4, pp. 1197–1210, 2009.

[21] C. Frithioff-Bøjsøe, M. A. Lund, U. Lausten-Thomsen, P. L. Hedley, O. Pedersen, M. Christiansen, J. L. Baker, T. Hansen, and J.-C. Holm, “Leptin, adiponectin, and their ratio as markers of insulin resistance and cardiometabolic risk in childhood obesity,” Pediatric diabetes, vol. 21, no. 2, pp. 194–202, 2020.

[22] G. Solinas, G. Germano, A. Mantovani, and P. Allavena, “Tumor-associated macrophages (tam) as major players of the cancer-related inflammation,” Journal of leukocyte biology, vol. 86, no. 5, pp. 1065–1073, 2009.

[23] M. Arnold, M. Leitzmann, H. Freisling, F. Bray, I. Romieu, A. Renehan, and I. Soerjomataram, “Obesity and cancer: an update of the global impact,” Cancer epidemiology, vol. 41, pp. 8–15, 2016.

[24] M. E. Williams, D. Howard, C. Donnelly, F. Izadi, J. G. Parra, M. Pugh, K. Edwards, K. Lutchman-Sigh, S. Jones, L. Margarit, et al., “Adipocyte derived exosomes promote cell invasion and challenge paclitaxel efficacy in ovarian cancer,” Cell Communication and Signaling, vol. 22, no. 1, p. 443, 2024.

[25] R. A. Ku-Carrillo, S. E. Delgadillo, and B. Chen-Charpentier, “A mathematical model for the effect of obesity on cancer growth and on the immune system response,” Applied Mathematical Modelling, vol. 40, no. 7-8, pp. 4908–4920, 2016.

[26] T. A. Yildiz, S. Arshad, and D. Baleanu, “New observations on optimal cancer treatments for a fractional tumor growth model with and without singular kernel,” Chaos, Solitons & Fractals, vol. 117, pp. 226– 239, 2018.

[27] N. Mohammad Mirzaei, N. Changizi, A. Asadpoure, S. Su, D. Sofia, Z. Tatarova, I. K. Zervantonakis, Y. H. Chang, and L. Shahriyari, “Investigating key cell types and molecules dynamics in pymt mice model of breast cancer through a mathematical model,” PLoS computational biology, vol. 18, no. 3, p. e1009953, 2022.

[28] M. P. Steinkamp, K. K. Winner, S. Davies, C. Muller, Y. Zhang, R. M. Hoffman, A. Shirinifard, M. Moses, Y. Jiang, and B. S. Wilson, “Ovarian tumor attachment, invasion, and vascularization reflect unique microenvironments in the peritoneum: insights from xenograft and mathematical models,” Frontiers in oncology, vol. 3, p. 97, 2013.

[29] C. Pouchol, Modelling interactions between tumour cells and supporting adipocytes in breast cancer. PhD thesis, UPMC, 2015.

[30] S. Oliver, M. Williams, M. K. Jolly, D. Gonzalez, and G. Powathil, “Exploring the role of emt in ovarian cancer progression using a multiscale mathematical model,” npj Systems Biology and Applications, vol. 11, no. 1, p. 36, 2025.

[31] R. M. Sutherland, “Cell and environment interactions in tumor microregions: the multicell spheroid model,” Science, vol. 240, no. 4849, pp. 177–184, 1988.

[32] A. Ghaffarizadeh, R. Heiland, S. H. Friedman, S. M. Mumenthaler, and P. Macklin, “Physicell: an open source physics-based cell simulator for 3-d multicellular systems,” PLoS computational biology, vol. 14, no. 2, p. e1005991, 2018.

[33] A. Ghaffarizadeh, S. H. Friedman, and P. Macklin, “Biofvm: an efficient, parallelized diffusive transport solver for 3-d biological simulations,” Bioinformatics, vol. 32, no. 8, pp. 1256–1258, 2016.

[34] M. Hoffmann, J. Gogola, and A. Ptak, “Adiponectin reverses the proliferative effects of estradiol and igf-1 in human epithelial ovarian cancer cells by downregulating the expression of their receptors,” Hormones and Cancer, vol. 9, pp. 166–174, 2018.

[35] G. M. Cooper, “The eukaryotic cell cycle,” National Library of Medicine, 2000.

[36] R. Sender and R. Milo, “The distribution of cellular turnover in the human body,” Nature medicine, vol. 27, no. 1, pp. 45–48, 2021.

[37] Y.-T. Chin, L.-M. Wang, M.-T. Hsieh, Y.-J. Shih, A. W. Nana, C. A. Changou, Y.-C. S. Yang, H.-C. Chiu, E. Fu, P. J. Davis, et al., “Leptin ob3 peptide suppresses leptin-induced signaling and progression in ovarian cancer cells,” Journal of Biomedical Science, vol. 24, pp. 1–13, 2017.

[38] F. Vikhanskaya, S. Vignati, P. Beccaglia, C. Ottoboni, P. Russo, M. D’Incalci, and M. Broggini, “Inactivation of p53 in a human ovarian cancer cell line increases the sensitivity to paclitaxel by inducing g2/m arrest and apoptosis,” Experimental cell research, vol. 241, no. 1, pp. 96–101, 1998.

[39] A. Brozovic, G. E. Duran, Y. C. Wang, E. B. Francisco, and B. I. Sikic, “The mir-200 family differentially regulates sensitivity to paclitaxel and carboplatin in human ovarian carcinoma ovcar-3 and mes-ov cells,” Molecular oncology, vol. 9, no. 8, pp. 1678–1693, 2015.

[40] S. Capellero, J. Erriquez, C. Battistini, R. Porporato, G. Scotto, F. Borella, M. F. Di Renzo, G. Valabrega, and M. Olivero, “Ovarian cancer cells in ascites form aggregates that display a hybrid epithelialmesenchymal phenotype and allows survival and proliferation of metastasizing cells,” International Journal of Molecular Sciences, vol. 23, no. 2, p. 833, 2022.

[41] M. A. Nieto, R. Y.-J. Huang, R. A. Jackson, and J. P. Thiery, “Emt: 2016,” Cell, vol. 166, no. 1, pp. 21–45, 2016.

[42] A. Bonnomet, L. Syne, A. Brysse, E. Feyereisen, E. Thompson, A. Noël, J.-M. Foidart, P. Birembaut, M. Polette, and C. Gilles, “A dynamic in vivo model of epithelial-to-mesenchymal transitions in circulating tumor cells and metastases of breast cancer,” Oncogene, vol. 31, no. 33, pp. 3741–3753, 2012.

[43] T.-T. Liao and M.-H. Yang, “Revisiting epithelial-mesenchymal transition in cancer metastasis: the connection between epithelial plasticity and stemness,” Molecular oncology, vol. 11, no. 7, pp. 792–804, 2017.

[44] J. Jiang, J. Li, X. Zhou, X. Zhao, B. Huang, and Y. Qin, “Exosomes regulate the epithelial–mesenchymal transition in cancer,” Frontiers in Oncology, vol. 12, p. 864980, 2022.

[45] C. B. Seymour and C. Mothersill, “Radiation-induced bystander effects—implications for cancer,” Nature Reviews Cancer, vol. 4, no. 2, pp. 158–164, 2004.

[46] T. K. Hei, H. Zhou, V. N. Ivanov, M. Hong, H. B. Lieberman, D. J. Brenner, S. A. Amundson, and C. R. Geard, “Mechanism of radiation-induced bystander effects: a unifying model,” Journal of Pharmacy and Pharmacology, vol. 60, no. 8, pp. 943–950, 2008.

[47] G. Nelson, J. Wordsworth, C. Wang, D. Jurk, C. Lawless, C. Martin-Ruiz, and T. von Zglinicki, “A senescent cell bystander effect: senescence-induced senescence,” Aging cell, vol. 11, no. 2, pp. 345–349, 2012.

[48] J. Maia, S. Caja, M. C. Strano Moraes, N. Couto, and B. Costa-Silva, “Exosome-based cell-cell communication in the tumor microenvironment,” Frontiers in cell and developmental biology, vol. 6, p. 18, 2018.

[49] J. P. Joseph, M. Harishankar, A. A. Pillai, and A. Devi, “Hypoxia induced emt: A review on the mechanism of tumor progression and metastasis in oscc,” Oral oncology, vol. 80, pp. 23–32, 2018.

[50] T. I. Croll, S. Gentz, K. Mueller, M. Davidson, A. J. O’Connor, G. W. Stevens, and J. J. Cooper-White, “Modelling oxygen diffusion and cell growth in a porous, vascularising scaffold for soft tissue engineering applications,” Chemical Engineering Science, vol. 60, no. 17, pp. 4924–4934, 2005.

[51] A. Moustakas and C.-H. Heldin, “Signaling networks guiding epithelial–mesenchymal transitions during embryogenesis and cancer progression,” Cancer science, vol. 98, no. 10, pp. 1512–1520, 2007.

[52] D. R. Grimes, A. G. Fletcher, and M. Partridge, “Oxygen consumption dynamics in steady-state tumour models,” Royal Society open science, vol. 1, no. 1, p. 140080, 2014.

[53] K. A. Landman and A. Q. Cai, “Cell proliferation and oxygen diffusion in a vascularising scaffold,” Bulletin of mathematical biology, vol. 69, pp. 2405–2428, 2007.

[54] Z. Huang and A. Xu, “Adipose extracellular vesicles in intercellular and inter-organ crosstalk in metabolic health and diseases,” Frontiers in immunology, vol. 12, p. 608680, 2021.

